# The architecture of resilience: a genome assembly of *Myrothamnus flabellifolia* sheds light on desiccation tolerance and sex determination

**DOI:** 10.1101/2025.06.02.657439

**Authors:** Rose A. Marks, John T. Lovell, Sarah B. Carey, Llewelyn Van Der Pas, Nyaradzai M. Chimukuche, Tomáš Brůna, Christopher Plott, Jenell Webber, Anna Lipzen, Juying Yan, Diane Bauer, Joanne Bentley, Jayson Talag, Kerrie Barry, Jane Grimwood, Jerry W. Jenkins, Jeremy Schmutz, Alex Harkess, Robert VanBuren, James Leebens-Mack, Jill M. Farrant

## Abstract

*Myrothamnus flabellifolia* is a dioecious resurrection plant endemic to southern Africa that has become an important model for desiccation tolerance. Here, we present a chromosome-level, haplotype-resolved reference genome for *M. flabellifolia* alongside transcriptomic profiling across a natural dehydration-rehydration cycle. The 1.28 Gb genome exhibits unusually consistent chromatin architecture with putative holocentric chromosome organization across highly divergent haplotypes. We identified an XY sexual system with a small sex-determining region on chromosome 8. Transcriptomic changes during dehydration varied with the severity of water loss and pointed to early suppression of growth, progressive activation of protective mechanisms, and reversible changes in transcript abundance upon rehydration. Co-expression networks and targeted gene family analyses revealed dynamic regulation of LEA and ELIP genes, with evidence of transcriptome “priming” in hydrated plants, with transcript abundance shifting toward highly disordered proteins during desiccation. Enrichment of ABA and light-responsive cis-regulatory elements in key genes support the existence of conserved stress response pathways. *M. flabellifolia’s* rich profile of phenolics and antioxidant genes highlight the overlap between desiccation tolerance and medicinal potential. Together, this study provides foundational resources for understanding the genomic architecture and reproductive biology of *M. flabellifolia* while offering new insights into the mechanisms of desiccation tolerance.

## INTRODUCTION

Plants have evolved diverse and sophisticated adaptations to survive drought. Understanding these adaptations and the mechanisms underlying them is important for addressing the challenges associated with water scarcity. Among these, desiccation tolerance—the ability to dry to a quiescent state and resume normal cellular function when rehydrated—is one of the most remarkable adaptations (Marks et al., 2025). Desiccation tolerance has played an important role in the evolution of terrestrial plants, enabling the colonization of land (Oliver et al., 2000, 2005; Farrant and Moore, 2011; Zhang et al., 2020) and eventually the domestication of crops through seed storage. Unlocking the molecular, cellular, and physiological mechanisms of desiccation tolerance could facilitate transformative innovations in agriculture, medicine, and material sciences, such as advanced xeropreservation techniques that enhance survival in dry conditions. As global droughts intensify, understanding these mechanisms is critical for safeguarding agricultural production, biodiversity conservation, and the communities that rely on these resources.

Desiccation tolerance has evolved repeatedly and convergently across diverse plant lineages (Oliver et al., 2000; VanBuren et al., 2019; Alejo-Jacuinde et al., 2020; Marks et al., 2024), but its mechanisms remain incompletely understood and overlap considerably with more generalized responses to water limitation in plants (Pardo et al., 2020). So called “resurrection plants” exemplify vegetative desiccation tolerance (Gaff, 1971; Gaff and Hallam, 1974; Griffiths et al., 2014; Tebele et al., 2021; Marks, 2024) and can survive months to years in a dry state, often under intense solar radiation and high heat, yet still resume full functionality within hours of rehydration (Oliver et al., 2020; Marks et al., 2021). This remarkable ability relies on complex molecular and physiological mechanisms that preserve cellular integrity and facilitate recovery (Giarola et al., 2017; Oliver et al., 2020; Gechev et al., 2021). Key components of these survival mechanisms include the accumulation of osmoprotectants, such as non-reducing sugars and polyols that stabilize proteins and membranes in a dry state (Oliver et al., 2011; Holzinger and Karsten, 2013; Vieira et al., 2017); the production of Late Embryogenesis Abundant (LEA) proteins (Hernández-Sánchez et al., 2022) and Early Light Induced Proteins (ELIPs) (VanBuren et al., 2019) that enhance protection and mitigate photooxidative damage; and the activation of antioxidant pathways that scavenge reactive oxygen species (ROS) (Dinakar et al., 2012).

Vegetative desiccation tolerance is also associated with the downregulation of photosynthesis and growth-related pathways (Challabathula et al., 2018), the modification of cell walls to increase flexibility (Moore et al., 2006, 2023; Neeragunda Shivaraj et al., 2018; Chen et al., 2019; Plancot et al., 2019; Alejo-Jacuinde et al., 2020), and the mobilization of proteostasis mechanisms, such as ubiquitination and autophagy (Zhu et al., 2015; Hibshman et al., 2023; VanBuren et al., 2023) that help to repair cellular components upon rehydration (Oliver et al., 2020). Taken together, it seems that dynamic changes in gene expression drive metabolic reprogramming, cellular protection, and repair processes that support survival during desiccation (Alejo-Jacuinde et al., 2020; Pardo et al., 2020; VanBuren et al., 2023; Marks et al., 2024; Zhang et al., 2024). Upon rehydration, these mechanisms are reversed or modified to support recovery and resume normal physiological functions (Oliver et al., 2020). Untangling the multifaceted and tightly coordinated responses to desiccation has proven challenging, but understanding of these processes would provide valuable insights into how life is maintained in a dry state.

*Myrothamnus flabellifolia* is one of the most iconic resurrection plants, having captured the attention of local communities and scientists for decades (Moore et al., 2007; Marks et al., 2022). It is locally abundant in some of the harshest environments in southern Africa, occupying a unique niche on rocky outcrops and cliffs. The species’s striking resilience is associated with the production of a suite of specialized compounds, including antioxidants, phenolics, and other diverse secondary metabolites, many of which have traditional and emerging medicinal significance (Bentley et al., 2019). Additionally, *M. flabellifolia* is dioecious, which provides a rare opportunity to study the evolution of sex determination and sexual dimorphisms in the context of extreme stress tolerance (Marks et al., 2022). Lastly, *Myrothamnus* (Myrothamnaceae) and *Gunnera* (Gunneraceae) are the only two extant genera within the order Gunnerales which is the sister lineage to the hyperdiverse Pentapetalae clade including all other core eudicots (Gunneridae) (Wanntorp et al., 2001; Cantino et al., 2007). These features position *M. flabellifolia* as a key system for exploring desiccation tolerance, evolutionary biology, and applications to human society.

Here, we present a chromosome-level, haplotype-resolved reference genome for *M. flabellifolia*, coupled with a high-resolution transcriptomic characterization of a natural dehydration and rehydration event in the field. By leveraging both genomic and transcriptomic resources, we explore the mechanisms underlying dioecy and sex determination, vegetative desiccation tolerance, and comment on the medicinal implications of the specialized secondary metabolism of *M. flabellifolia*.

## RESULTS

### Study organism and accessions

*M. flabellifolia* is an iconic resurrection plant in the eudicot lineage Gunnerales. Myrothamnaceae contains only one genus with just two species in it; *M. flabellifolia* and *M. moschatus* (Moore et al., 2007). *M. moschatus* is endemic to Madagascar while *M. flabellifolia* is distributed throughout continental southern Africa in disjunct populations (Figure 1A). Both species occupy a narrow ecological niche, restricted to rocky sites, with minimal soil, and intense abiotic stresses (Marks et al., 2022; Wan et al., 2024). In addition to being desiccation tolerant, *M. flabellifolia* is dioecious with separate male and female individuals (Figure 1B-E). *M. flabellifolia* also produces a robust profile of secondary compounds with important traditional and emerging medicinal applications (Bentley et al., 2019). For the current study, *M. flabellifolia* plants were collected from three populations across the species range (Figure 1A). A genome assembly was generated for a single heterogametic (male) individual from the selected focal site, Swebe Swebe Nature Reserve in Limpopo, South Africa, and transcript profiles were generated for 12 individuals during a natural dehydration-rehydration event (see ***Desiccation tolerance*** section below) at Swebe Swebe Nature Reserve.

**Figure 1.**
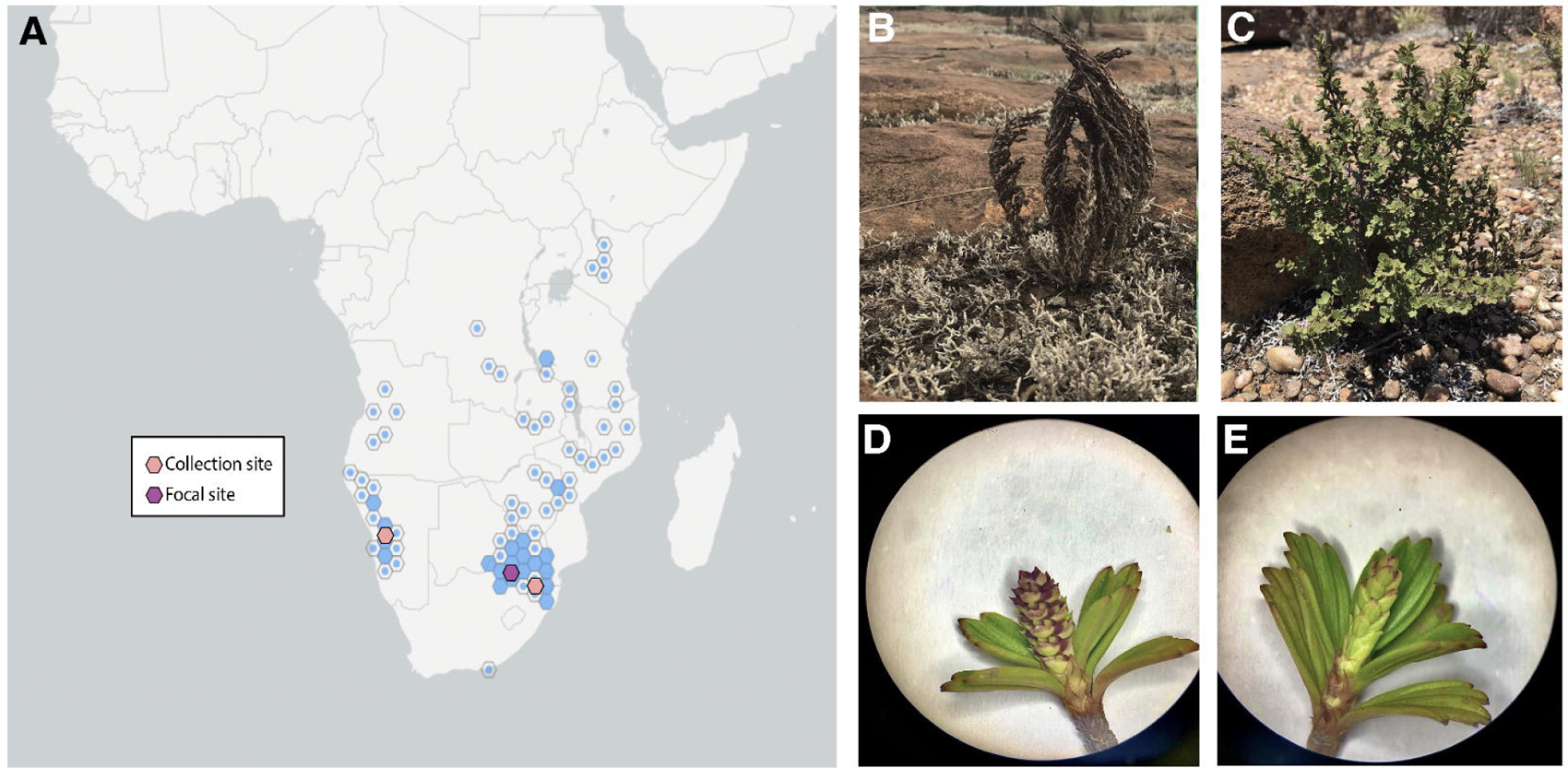
Distribution and anatomy of *Myrothamnus flabellifolia*. (A) *Myrothmnus flabellifolia* is distributed across southern Africa in disjunct populations. Plants for the current study were collected from three sites–two in South Africa and one in Namibia. A single focal site was selected for detailed investigations, in Limpopo, South Africa. *Myrothmnus flabellifolia* is shown in both (B) desiccated and (C) hydrated conditions in its native habitat. Floral morphology of (D) male and (E) female plants.

### The unusual genome structure of M. flabellifolia

We generated a chromosome-level, haplotype-resolved genome assembly for *M. flabellifolia* using deep PacBio HiFi sequencing technology (69.05x per haplotype; mean read length = 15,920bp). The final genome assembly, which was scaffolded with 61.5x HiC and polished with 54x Illumina reads, spans ∼1.28 Gb per haplotype, with Haplotype 1 (HAP1) built from 195 contigs (N_50_ = 11.4Mb) and HAP2 from 169 contigs (N_50_ = 13.7Mb, Table 1). The assemblies were annotated with gene models supported by evidence derived from 12 RNA-seq libraries (>909M total reads) constructed using RNAs extracted from tissues collected across five conditions and 12 Iso-seq libraries constructed from pooled RNAs (>15M reads). The resulting annotation included 22,809/22,922 (HAP1/HAP2) protein-coding gene models both with perfect Eukaryote and ≥ 99% Embryophyta BUSCO scores.

**Table 1:** Genome assembly and gene annotation statistics. Statistics are reported by haplotype, but phasing was done individually for each chromosome.

|  | Haplotype 1 | Haplotype 2 |
| --- | --- | --- |
| **Chromosome Sequence** | 1,283.3 Mb | 1,273.1 Mb |
| **Contig total** | 195 | 169 |
| **Scaffold $N_{50}$** | 134.5 Mb | 133.8 Mb |
| **Contig $N_{50}$** | 11.4 Mb | 13.7 Mb |
| **Number genes / transcripts** | 22,809 / 47,534 | 22,922 / 47,563 |
| **Annotation BUSCO completeness**<br>**(embryophyta_odb10/eudicots_odb10)** | 99.0 / 98.7 % | 99.1 / 98.5 |
| **Annotation PSAURON** | 93.5 | 93.5 |

Comparisons of homologous chromosome assemblies between the two haplotypes reveal a high level of nucleotide diversity with an average of 100.58 SNPs per 10kb. We also observed 63 (45.5Mb total) large insertions, deletions, or inversions >100kb. Of particular note was a large inversion on Chr01 spanning 18.4Mbp (11.3% of the chromosome). Commensurate with this molecular variation, we observed significant gene content variation between the haplotypes: of the 22,902 total gene families, 2,605 (11.3%) and 1,254 (5.5%) exhibit presence-absence or copy number variation respectively.

Despite the degree of nucleotide and structural variation between homologous chromosomes, gene and repeat density is remarkably uniform across the *M. flabellifolia* genome. Every 10Mb genomic window (5Mb overlapping) of HAP1 was annotated with 0.5-1.5Mb (5-15%) of genic sequence and 60-82% of EDTA-defined repeat sequences. Visually, this consistency appears to be an outlier among flowering plants and more in line with expectations from Bryophytes (Healey et al., 2023). Indeed, across 1000 non-overlapping windows the % of gene density standard deviation in both *M. flabellifolia* (SD HAP1 = 3.6%, HAP2 = 3.8%) was closer to that of *Sphagnum fallax* (7.3%) than the highly variable eudicot genome structures like those of *Arabidopsis thaliana* (23.0%), *Mimulus guttatus* (21.7%) or cotton (10.6%), all of which retain the majority of their genes in repeat-poor chromosome arms, and have few genes in the highly repetitive pericentromeric regions that surround a readily identifiable tandem-repeat filled centromere. However, no such structural variation was apparent in *M. flabellifolia*. Not only were we unable to identify pericentromeric repetitive regions, but none of the three major tools to computationally identify centromeres could find clusters of putative centromeric repeats. When combined with a remarkably consistent 3D chromatin architecture visible in genome-wide HiC data, these lines of evidence potentiate a holocentric chromosome organization. However, further molecular and visual validation of holocentric chromosomes will be required to confirm this hypothesis.

### Sexual dimorphisms and identification of the sex determining region

Both species in *Myrothamnus* are dioecious with distinct male and female anatomy, as are some members of *Gunnera,* suggesting there could be a shared origin of dioecy prior to the divergence of these genera. However, it is currently unknown what the sex chromosome system is in these species, nor what their contribution is to sexually dimorphic traits. In *M. flabellifolia*, inflorescences of both males and females consist of densely packed florets in a simple arrangement with extremely short pedicles, but male inflorescences have bracts and female inflorescences are bract-less (Figure 1). The male flowers develop three to six stamens that dehisce longitudinally, while females develop three basally attached carpels. Floral organs are desiccation tolerant in both sexes, but male flowers become desiccation sensitive after pollen dehiscence (Moore et al., 2007).

Illumina DNA sequencing data of 10 accessions from across the collection sites (Figure 1A) were used to predict the sex determination system. To first determine whether females or males contain the heterogametic sex chromosome pair in *M. flabellifolia*, we used a *k*-mer based analysis (Carey et al., 2024a, b). We found four times more male-specific *k*-mers than female-specific, indicating an XY sex-determination system. To identify the Y chromosome in the genome assembly, and delimit the boundary of the sex-determination region (SDR) from the recombining pseudoautosomal region (PAR), we mapped the putative Y-specific *k*-mers to both haplotypes. We found the SDR on the Y chromosome to be located on chromosome 8 of HAP2 spanning only ∼700 Kb (Figure 2), making it among some of the smaller SDRs identified in plants, similar in size to some poplars (Renner and Müller 2021). Despite the smaller size, there are clear structural variants between the SDR and the homologous region to the X (Figure 2).

**Figure 2.**
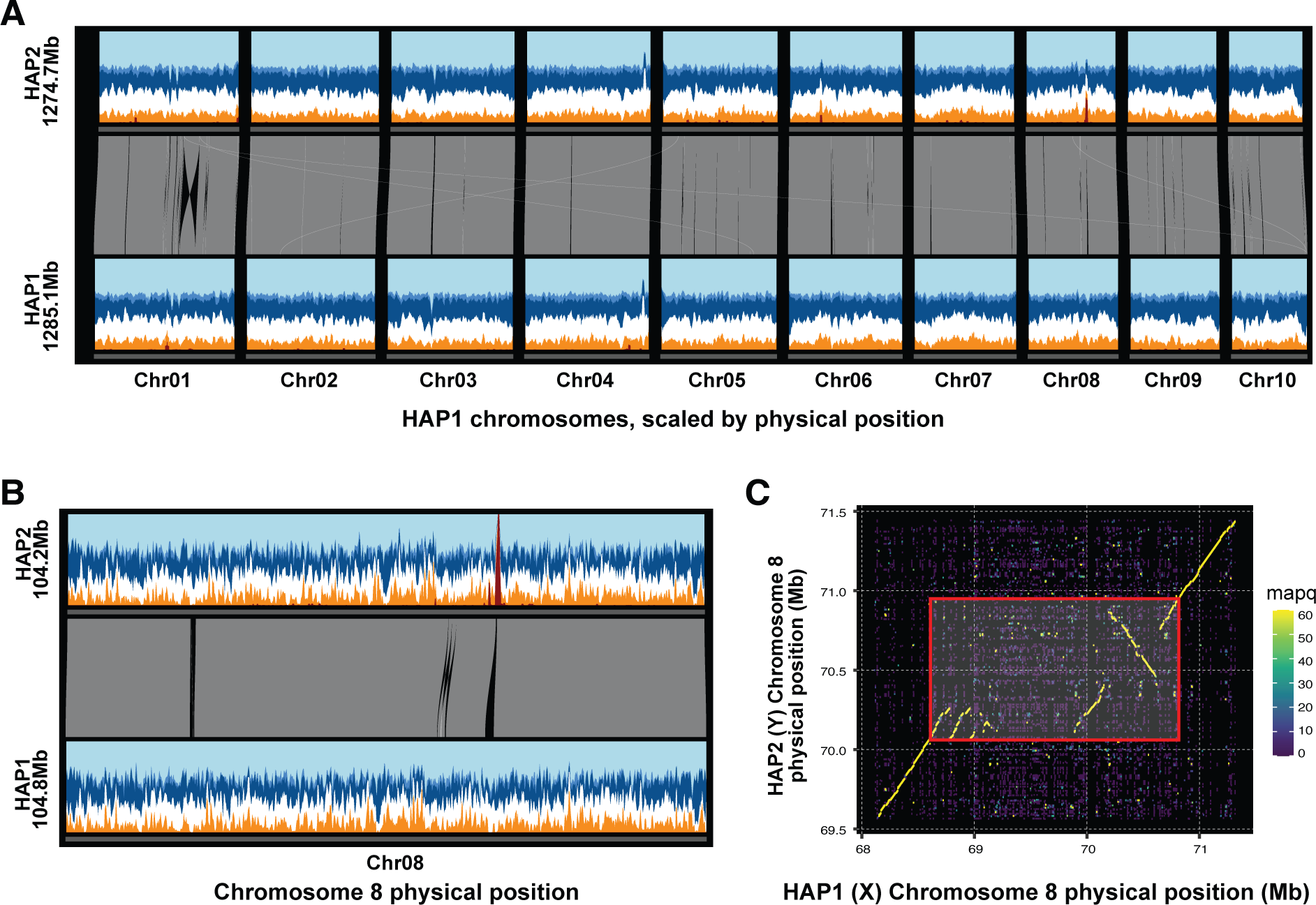
The unusual genome structure of *Myrothamnus flabellifolia*. (A) Genome-wide synteny (grey links) show that one large (Chr01) and several small inversions exist between the two haplotypes of the reference genome, but otherwise the vast majority of the sequences are syntenic and readily alignable. In contrast to typical angiosperm genomes, repeat (Ty3: dark blue, Ty1/Copia: medium blue, other EDTA-annotated repeat: light blue) and gene (orange) density is remarkably consistent without an obvious repeat-dense pericentromere surrounding putative centromeres. Gene, repeat and male-specific kmer (dark red) contributions were calculated within 100kb-overlapping 2.5Mb sliding windows. (B) This structure is further illustrated in Chromosome 8 where the sex determining region occurs, as indicated by a peak of male-specific kmers (dark red peak on hap 2) and a conspicuous presence absence variant. Sliding window method follows that of panel (A) except with smaller 250kb widths. (C) Further zooming in on the SDR (red box) and a 500kb buffer reveals that the sequence insertion into the X (HAP1) sequence occurs directly in the center of the SDR and is bounded by a partially degraded 5’ sequence triplication and a 3’ large inversion/duplication. Points in the dotplot are 50bp-overlapping 200bp windows of HAP1 mapped to the HAP2 sequence. Mapq is the minimap2-defined mapping quality of these hits. Sliding windows, synteny maps and dotplots were calculated and plotted with DEEPSPACE (<u>github.com/jtlovell/DEEPSPACE</u>).

Interestingly, the homologous region to the X is larger than the SDR of the Y as a consequence of tandem triplication of one end of the region and accumulation of repetitive elements comprising a nearly 750 kb X-specific segment. Whereas the Y-specific SDR contains just six gene models, we identified 11 X-linked homologs comparing the two haplotypes in an OrthoFinder v2.5.2 analysis (Emms and Kelly, 2015, 2019). One single copy gene on the Y, with four orthologous copies within the 5’ triplicated region on the X (Figure 2) was predicted to encode a Kinesin-1 like protein. A homolog in rice has been shown to be important for anther dehiscence and male meiosis (Zhou et al., 2011), suggesting it may have a role in male development, but the Arabidopsis homolog, *AtKin-1*, influences female gametophyte development (Wang et al. 2014). The identification of the XY chromosomes, and gene content in the SDR and homologous X-region, provides useful information for investigating and predicting the genetic control of dioecy across the flowering plant phylogeny <u>(Carey et al. 2021)</u>. For example, future work will assess a role for Kinesin-1 homologs in sex determination in *M. flabellifolia* and other dioecious plants.

Interestingly, no sexual dimorphisms in desiccation tolerance have been described in *M. flabellifolia*, but population-level analyses have identified male-biased sex ratios in more arid environments (including the focal site) compared to female dominance in mesic regions (Marks et al., 2022). These patterns are likely driven by dimorphic resource allocation, with females requiring more carbon and water for seed maturation and males allocating nitrogen and other resources for inflorescence and pollen production. Secondary sexual dimorphisms, including faster growth in females and greater inflorescence production in males have also been identified and likely influence population dynamics across environmental gradients (Marks et al., 2022).

### Dynamic responses to desiccation and rehydration

To better understand the molecular response to desiccation in *M. flabellifolia,* we performed high-resolution transcriptomic profiling 12 individual *M. flabellifolia* plants across a six-day natural dehydration-rehydration time course in the field at Swebe Swebe Nature Reserve in Limpopo, South Africa. The plants exhibited variable rates of drying based on water loss, likely due to differences in micro-niche, and we classified them into three groups based on the minimum relative water content (RWC) they reached during the time course. Some plants desiccated rapidly and completely, with RWC dropping below 10%, and we classified these as the “desiccation” group. Others experienced slower but still severe dehydration, reaching 10– 30% RWC, and were classified as “severe dehydration”. A third group underwent only mild dehydration, with RWC declining to 60–85% and were classified as “mild dehydration” (Figure 3). This stratification corresponds to the early and late stages of dehydration proposed to characterize drying in resurrection plants (Farrant and Hilhorst, 2021). These drying groups were leveraged for statistical comparisons between mild dehydration, severe dehydration, and full desiccation.

**Figure 3.**
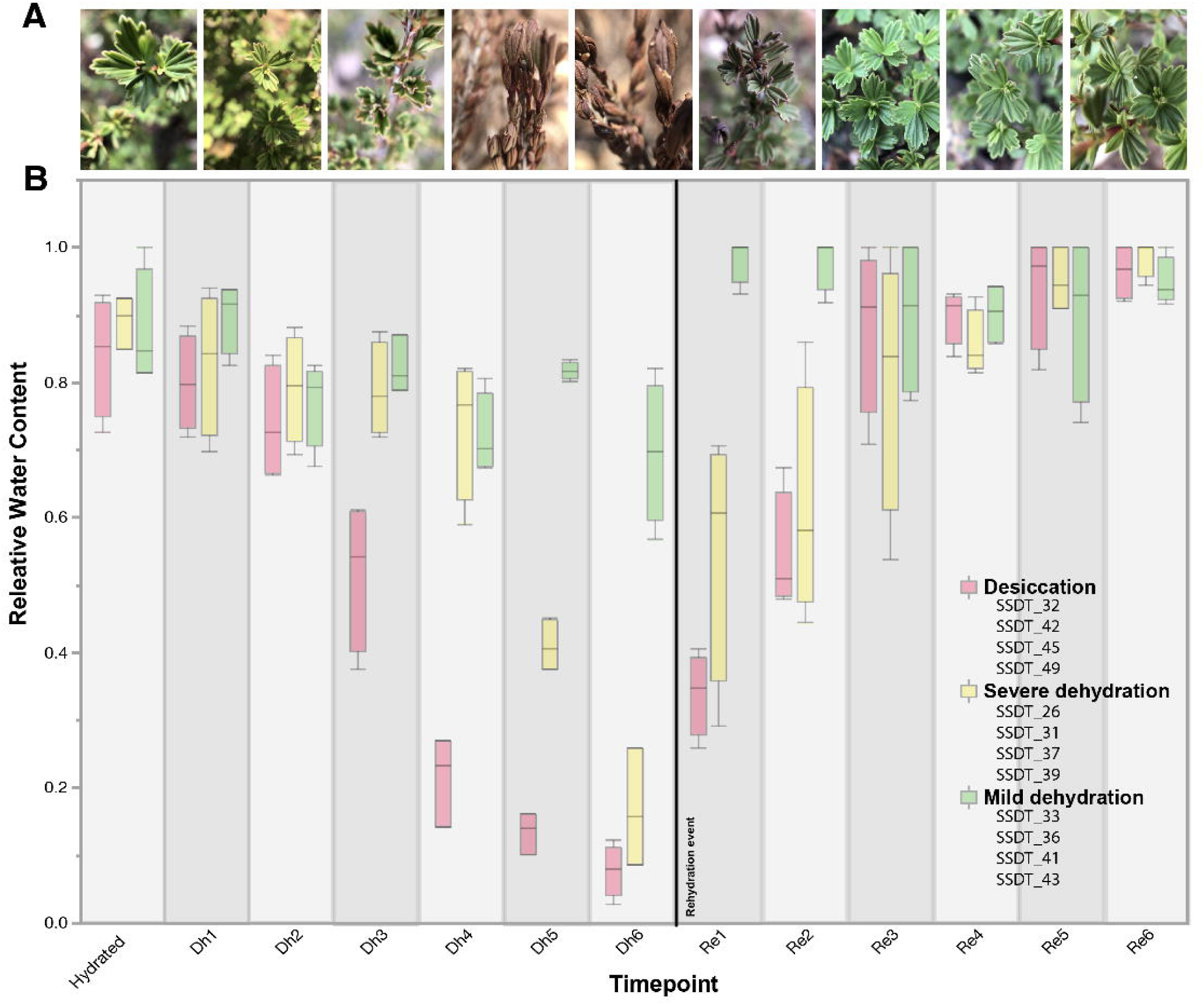
Drying dynamics of *Myrothamnus flabellifolia*. (A) Representative photographs of plants at different stages of dehydration and rehydration. (B) Relative water content (RWC) of plants throughout the 6-day dehydration-rehydration time course. Plant genotypes (e.g., SSDT_26) are listed and grouped / colored by their drying groups.

### Overview of broad patterns via dimension reduction analyses

To visualize broad changes in transcript abundance across the dehydration-rehydration time course, we performed principal component analysis (PCA). PCA revealed a strong association between transcript abundance and RWC, with the first principal component (PC1) explaining 70% of the total variation in transcript abundance and separating samples primarily on water status (Figure 4A). We also conducted targeted PCA for each drying group separately. These analyses identified increasing segregation of samples under more severe drying conditions, emphasizing an escalating transcriptomic response to progressive dehydration (Figure 4A).

**Figure 4.**
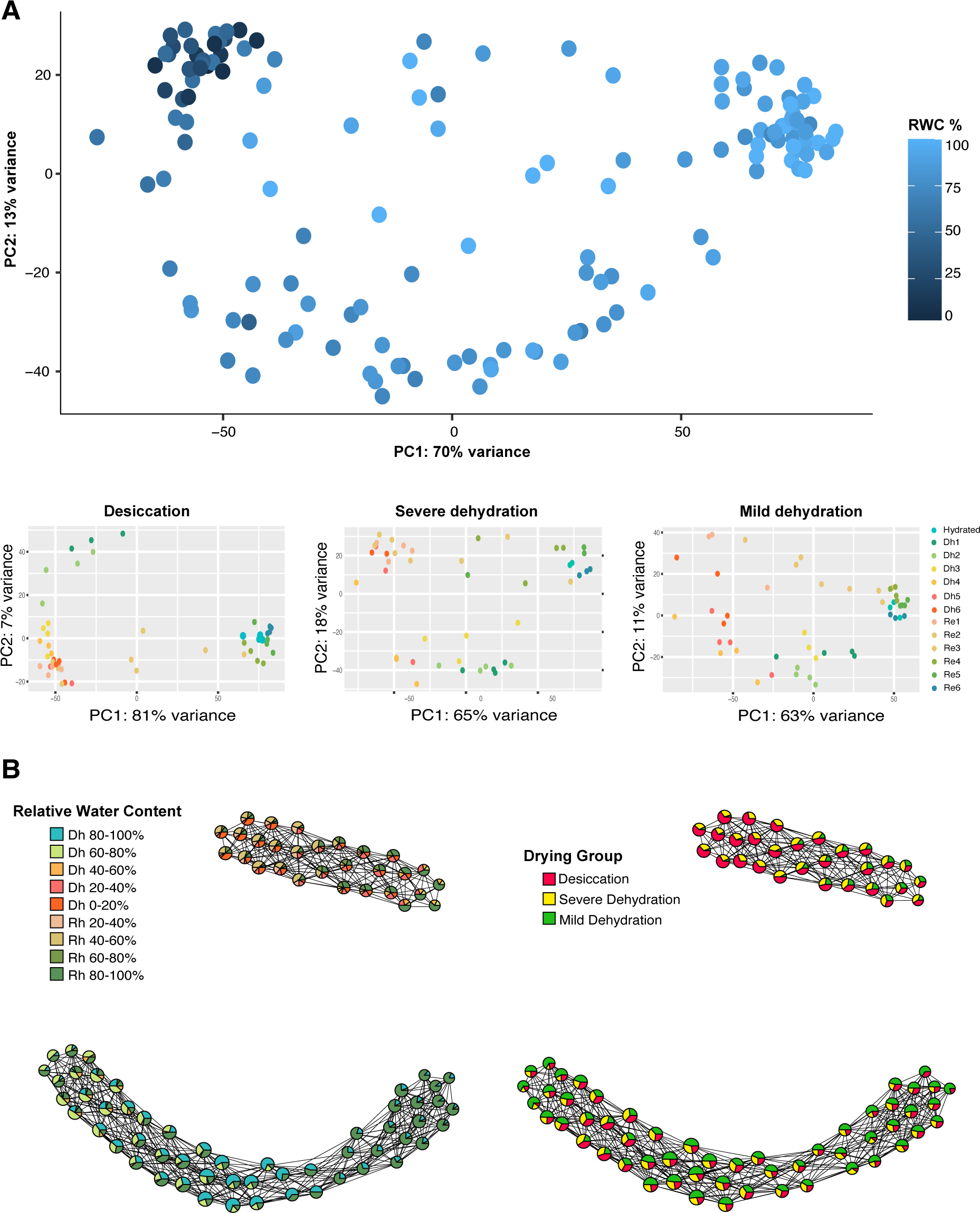
Gene expression during the dehydration time course. (A) PCA of transcript abundance for all plants colored by relative water content. PCAs of transcript abundance in different drying groups, colored by sampling time. (B) Topological data analyses of transcript abundance. Each node on the graph represents a cluster of similar RNAseq samples, and the node color depicts the identity of samples within that cluster. Connections between nodes signify shared samples among the intersecting clusters.

Although PCA provided some dimensionality reduction, residual variability—arising from heterogeneity, experimental noise, or genotype-level differences in the data—may have masked underlying biological patterns. To address this, we applied topological data analysis (TDA) using the Mapper algorithm <u>(van Veen et al. 2019; Palande et al. 2023; Marks et al. 2024)</u>, which provides a flexible and scalable approach for exploring high-dimensional, sparse datasets.

Mapper requires a lens function, a user-defined feature that shapes how the data are clustered and connected. We used RWC as the lens, anchoring it to the fully hydrated condition, to reflect physiological water status across samples. This allowed us to explore expression changes along a continuum of dehydration and recovery. The resulting Mapper graph showed that water status was the primary driver of expression variation, followed by the imposed drying group classification (Figure 4B). We found no clear patterns related to the sex or genotype of accessions, indicating that transcriptomic changes during desiccation and rehydration are predicted primarily by water status and drying severity, with minimal differences across individuals, genotypes, or sexes.

### Differentially abundant transcripts showcase escalating response to desiccation

Next, we quantified changes in transcript abundance throughout the dehydration-rehydration time course by identifying differentially abundant transcripts (DATs). DATs were identified separately for each drying group by comparing transcript abundance at each sampling time point to baseline hydrated conditions. This approach allowed us to differentiate between responses associated with progressive water loss. Our results revealed an escalation in the number of DATs as dehydration intensified throughout the time course (Figure 5A). Upon rehydration, transcript abundance largely returned to baseline levels, highlighting the reversible nature of these molecular changes (Figure 5A). Not surprisingly, changes in transcript abundance were more pronounced in plants that underwent full desiccation compared to those that only experienced mild dehydration (Figure 5A). In plants that underwent full desiccation more than 60% of the transcripts exhibited significant changes in abundance, but even under mild dehydration more than 40% of the transcripts exhibited differential abundance, reflecting the early induction of massive transcriptomic remodeling associated with drying.

**Figure 5.**
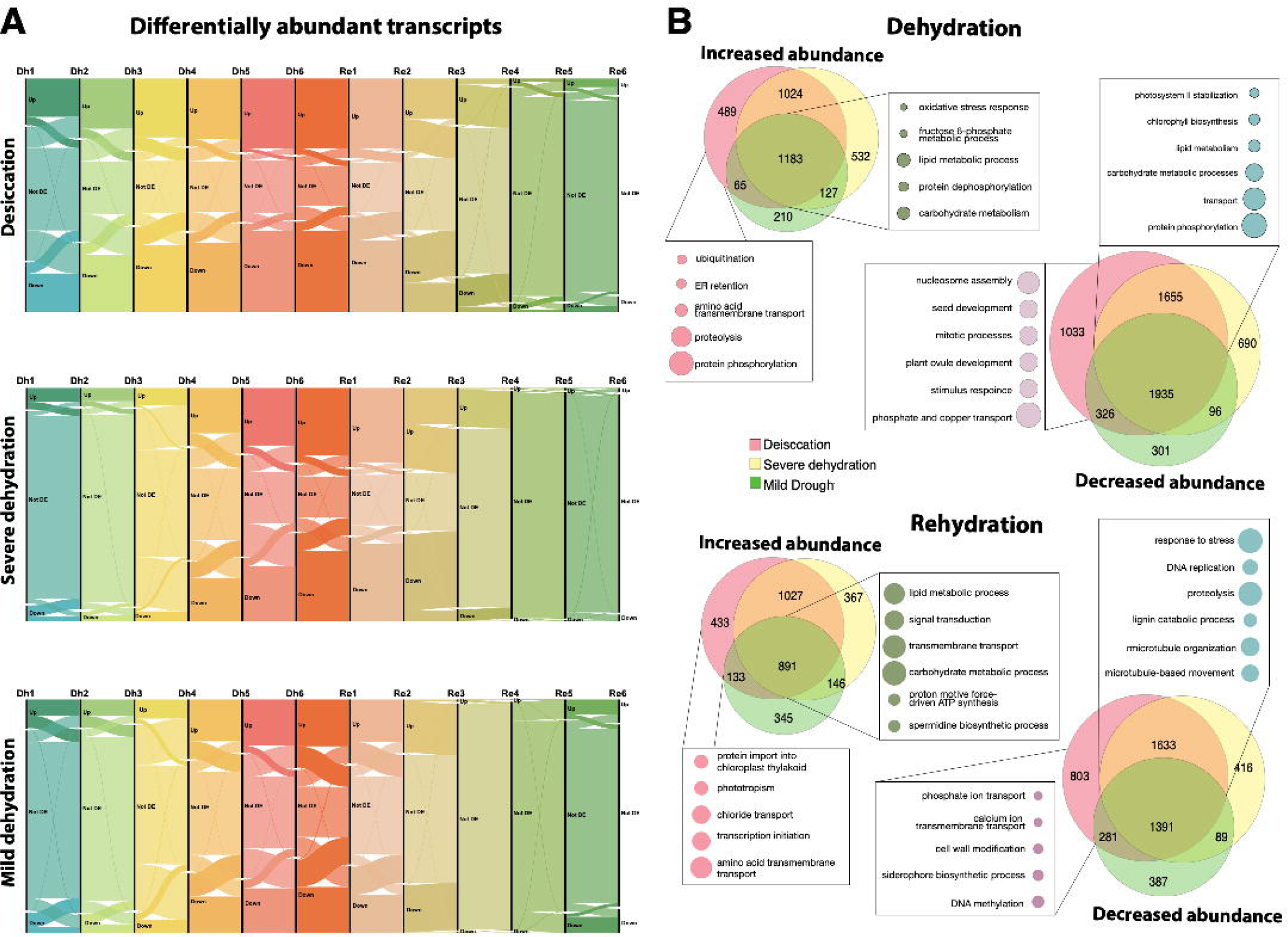
Differentially abundant transcripts. (A) Alluvial diagrams showing the number and flow of differentially expressed genes at each time point for each of the three drying groups. (B) Venn diagrams showing the number of overlapping and unique DATs across the three drying groups for up and down regulated genes along with a selection of the prominent enriched gene ontology categories that were shared among all plants vs those that were unique to the most extreme stages of desiccation.

To capture the key transcriptomic shifts in each drying group, we generated non-redundant lists of DATs that either increased or decreased in abundance during dehydration and separately during rehydration. Comparing these lists across the three drying groups revealed both shared and unique changes in transcript abundance (Figure 5B). Most DATs were common to all drying groups, reflecting the early activation of water deficit responses initiated by all plants. However, some unique DATs were identified only during the later stages of desiccation. These DATs likely represent responses to, and consequences of, the intense stress of desiccation, and provide valuable insights into the progressive activation and repression of molecular pathways during desiccation.

We conducted gene ontology (GO) enrichment analysis of shared and unique DATs to identify key pathways affected by dehydration and rehydration (Figure 5B). During drying, all plants exhibited a decrease of photosynthesis-related processes, including photosystem II stabilization and chlorophyll biosynthesis, alongside fatty acid biosynthesis, lipid metabolism, protein phosphorylation, DNA replication, and cell wall modification (e.g., xyloglucan metabolism). These changes likely reflect an overall energy-saving strategy to minimize metabolic demands, ROS production, and damage under water stress. In plants experiencing mild and severe dehydration, circadian rhythm regulation, auxin responses, and glutamine biosynthesis were suppressed, suggesting a shift away from growth to prioritize survival under water deficit conditions. Plants that experienced full desiccation displayed unique reductions in cell division, mitotic processes, ovule development, stimulus response, as well as phosphate and copper transport, indicating a more complete halt in growth and resource redistribution (Figure 5B).

Pathways related to oxidative and salt stress increased in all plants during drying. All plants also exhibited an increase in transcripts related to metabolic and catabolic processes of carbohydrate metabolism (e.g., sucrose transport, glycolysis, glyoxylate cycle, and malate metabolism). These changes likely enhance energy production, osmoprotection, and cellular defense against stress. During severe dehydration, transcripts related to spermidine and spermine biosynthesis, vitamin E, and terpenoid biosynthesis were enriched, which are likely involved in cellular protection against oxidative damage. RNA regulation, nucleotide biosynthesis, and RNA surveillance were also abundant during more extreme dehydration indicating changes in nucleic acid regulation. In plants experiencing complete desiccation, protein phosphorylation, ubiquitination, ER retention, and proteolysis pathways were enriched, suggesting that mechanisms to manage protein stability, recycle damaged proteins, and prevent damage are important during final stages of desiccation (Figure 5B).

During rehydration, all plants exhibited continued suppression of DNA replication, microtubule organization, and cell wall-related processes (e.g., cellulose and microfibril organization), suggesting a delay in resuming growth until environmental conditions stabilize and damage is repaired. Processes related to cell growth, phototropism, and flowering also remained suppressed, indicating a focus on repair and recovery prior to regeneration. Plants experiencing complete desiccation showed unique reduction in phosphate transport, calcium transport, lipid metabolism, and DNA methylation during rehydration, possibly reflecting delayed resumption of metabolic balance and long term expression remodeling via epigenetic marks (Figure 5B).

Many transcripts increased during rehydration in all plants, including those related to spermine and spermidine biosynthesis, ethanol oxidation, sulfate assimilation, terpenoid biosynthesis, and RNA surveillance, suggesting a balance between continued stress response and metabolic recovery. Some pathways were uniquely increased in plants that experienced full desiccation, including autophagy, ribosomal subunit biogenesis, phototropism, and chloride transport, pointing towards extensive repair mechanisms needed to restore normal cellular function after desiccation (Figure 5B). Taken together, these patterns highlight the interplay between preemptive suppression of growth and development and activation of protective mechanisms required for desiccation tolerance. The ability to halt growth while upregulating repair and defense pathways is likely critical to survival and recovery.

### Modules of co-expressed genes exhibit dynamic activation and repression

To further explore changes in transcript abundance across the dehydration-rehydration time course, we identified modules of co-expressed genes and plotted the expression of these modules across the time course for each of the three drying groups (Figure 6). Many of the modules exhibited similar expression patterns in all of the drying groups (e.g., black, cyan, and magenta modules; Figure 6 left-hand column of panels) and these likely represent core responses to water loss established in the early stages of drying. These modules were enriched for GO terms involved in carbohydrate related processes such as gluconeogenesis, glycolysis, TCA cycle, sucrose, glucose, and fructose-6-phosphate metabolism, cell wall biogenesis and modification, cellulose microfibril organization and biosynthesis, and lignin catabolism. These findings align with the DAT results, which highlighted carbohydrate-related processes and cell wall modifications as early induced responses to dehydration. These modules contained other hallmarks of desiccation tolerance, such as trehalose biosynthesis, superoxide metabolism, glutathione and glutamine biosynthesis, and GABA catabolism, indicating that protective mechanisms against oxidative damage and metabolic adjustments are also initiated early, even before plants experience full desiccation.

**Figure 6.**
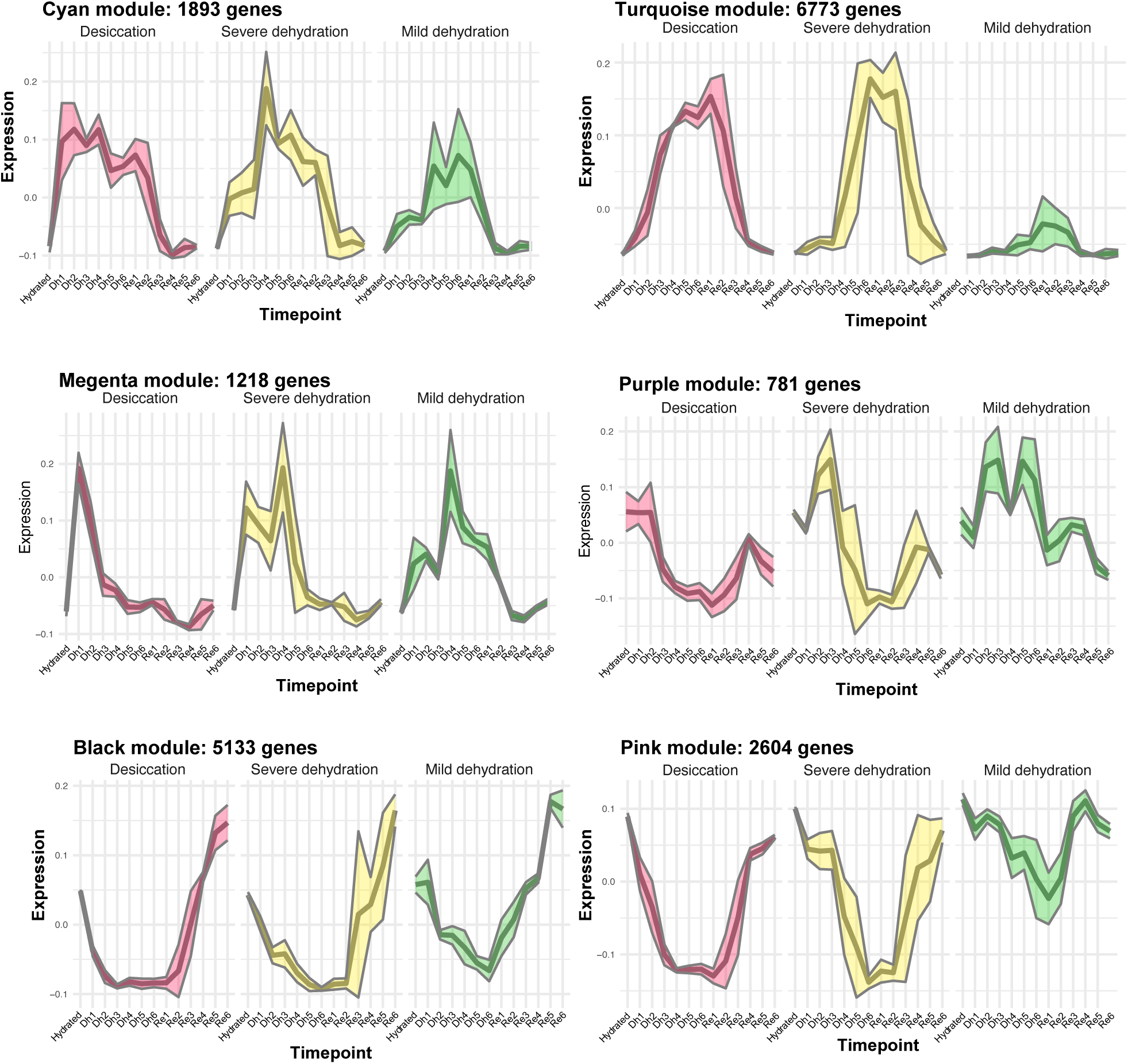
Expression profiles of co-expressed modules in each of the three drying groups. Modules in the left column have similar expression in all drying groups whereas modules in the right column show contrasting expression across different drying groups.

In contrast to the broadly conserved modules, certain key modules (e.g., pink, purple, and turquoise modules; Figure 6 right-hand column of panels) displayed distinct expression patterns related to the severity of dehydration. The pink module, for example, was downregulated only in plants experiencing the severe dehydration and desiccation but remained near baseline levels under mild dehydration. This module was functionally enriched for nucleoside triphosphate biosynthesis, microtubule and cytoskeleton-related processes, and amino acid-related pathways such as tryptophan biosynthesis. These results indicate that processes related to cell division and mitotic organization were increasingly downregulated as plants reached full desiccation.

The suppression of these processes likely reflects a halt in energy-intensive cellular activities to conserve resources for survival. The turquoise module also showed contrasting patterns between drying groups, with increased transcript abundance only in severe dehydration and desiccation, and minimal activation in mild dehydration conditions. This module was enriched for GO terms associated with nucleic acid regulation, including telomere maintenance, chromatin remodeling, DNA-templated transcription, mRNA splicing, and amino-tRNA aminoacylation, suggesting that regulation of transcription and translation is critical in later stages of drying. The turquoise module also included terms related to protein homeostasis, such as proteolysis and protein trafficking, paralleling DAT results that indicated a similar increase of protein phosphorylation and ubiquitination during more extreme stages of dehydration. Together, these findings suggest that plants experiencing severe dehydration activate specialized pathways for genomic stability, transcriptional regulation, and protein quality control that may be key for survival under extreme water stress. The purple module was uniquely activated in plants experiencing mild dehydration but suppressed in plants experiencing more extreme dehydration and desiccation. This module was enriched for circadian regulation, phototropism, fatty acid beta-oxidation, protein secretion, urea catabolism, spliceosomal snRNP assembly, and rRNA modification. The selective activation of this module in mild dehydration suggests that these pathways may represent early-stage responses aimed at maintaining homeostasis under less severe stress while also initiating basic preparation for increasing dehydration. The downregulation of these processes in more extreme conditions may reflect a shift from maintaining normal cellular function towards a committed prioritization of survival mechanisms.

### Antioxidant capacity and implications for medicine

The resilience of *M. flabellifolia* to desiccation can be partly attributed to its rich composition of phenolic compounds (Bentley et al., 2019), which play an important role in protecting the plant against oxidative stress and have intriguing medicinal properties. Flavonoids, hydroxycinnamic acids, and anthocyanins, among others, contribute significantly to the antioxidant capacity of *M. flabellifolia* by acting as free-radical scavengers targeting ROS, such as superoxide radicals, which are generated during periods of stress. These phenolic compounds help maintain the structural integrity of plant tissues and protect cells from oxidative damage (Kranner et al., 2002). The same antioxidant and stress-mitigating properties that confer desiccation tolerance are also key to the medicinal relevance of *M. flabellifolia*, as many of these compounds exhibit pharmacological potential, including anti-inflammatory and neuroprotective effects.

Several GO terms linked to radical scavenging activity and stress responses were identified across the co-expression modules activated during dehydration. In the turquoise coexpression module, which peaked most dramatically under both desiccation and severe dehydration (Figure 6), several GO terms were closely linked to redox homeostasis (e.g., superoxide metabolic process, glutathione biosynthetic process), metabolic pathways (e.g., TCA cycle and glycolytic process), and signaling pathways (e.g., intracellular signal transduction). The abundance of phenolic compounds in desiccated *M. flabellifolia* tissues (Bentley et al., 2019) aligns with these transcriptomic responses. Specifically, gallic acid and its galloyl derivatives, ellagic acid derivatives, various quercetin derivatives, and anthocyanins (such as delphinidin and cyanidin glycosides) are strongly associated with superoxide metabolic processes, while naringenin, quercetin derivatives, and anthocyanins modulate intracellular signal transduction pathways, thereby influencing cellular stress responses and potentially contributing to their medicinal efficacy. These functions, while crucial for desiccation tolerance, may also be relevant for biomedical applications, particularly in understanding how antioxidant compounds can contribute to genome stability and cellular repair mechanisms in other biological systems.

*Dynamic changes in the abundance of key genes highlight complex regulatory networks* Both the Late Embryogenesis Abundant (LEA) and Early Light-Induced Protein (ELIP) gene families have well documented roles in desiccation tolerance (VanBuren et al., 2019; Hernández-Sánchez et al., 2022), and we conducted targeted analyses to examine changes in the abundance of LEA and ELIP transcripts in *M. flabellifolia* across the dehydration-rehydration time course. LEAs have long been recognized as key players in stress response and are generally regarded as molecular protectants thought to stabilize proteins and membranes during water loss. LEAs typically have a high degree of intrinsic disorder (Hernández-Sánchez et al., 2022) and have roles in stabilizing the bioglasses generated during desiccation (Rascio and Rocca, 2005). ELIPs, on the other hand, have emerged more recently as critical players in desiccation tolerance, with a presumed role in preventing photooxidative damage through their chlorophyll-binding activities (Hutin et al., 2003). ELIPs are significantly expanded in the genomes of all sequenced resurrection plants and typically occur in tandem arrays (VanBuren et al., 2019; Marks et al., 2024).

Both LEAs and ELIPs are among the most highly expressed transcripts during desiccation across numerous species, and *M. flabellifolia* is no exception. Most *M. flabellifolia* LEA and ELIP transcripts increased dramatically in abundance during dehydration. Interestingly, a few specific LEA and ELIP transcripts showed the opposite pattern and were abundant during hydrated conditions but declined as dehydration progressed. This pattern suggests a baseline level of protection or “priming” in *M. flabellifolia,* that may equip the plants for upcoming rapid water loss. As dehydration progressed, expression shifted from one set of LEAs and ELIPs to a different subset, reflecting a possible transition from general protective mechanisms to those optimized specifically for dehydration and desiccation (Figure 7A and B).

**Figure 7.**
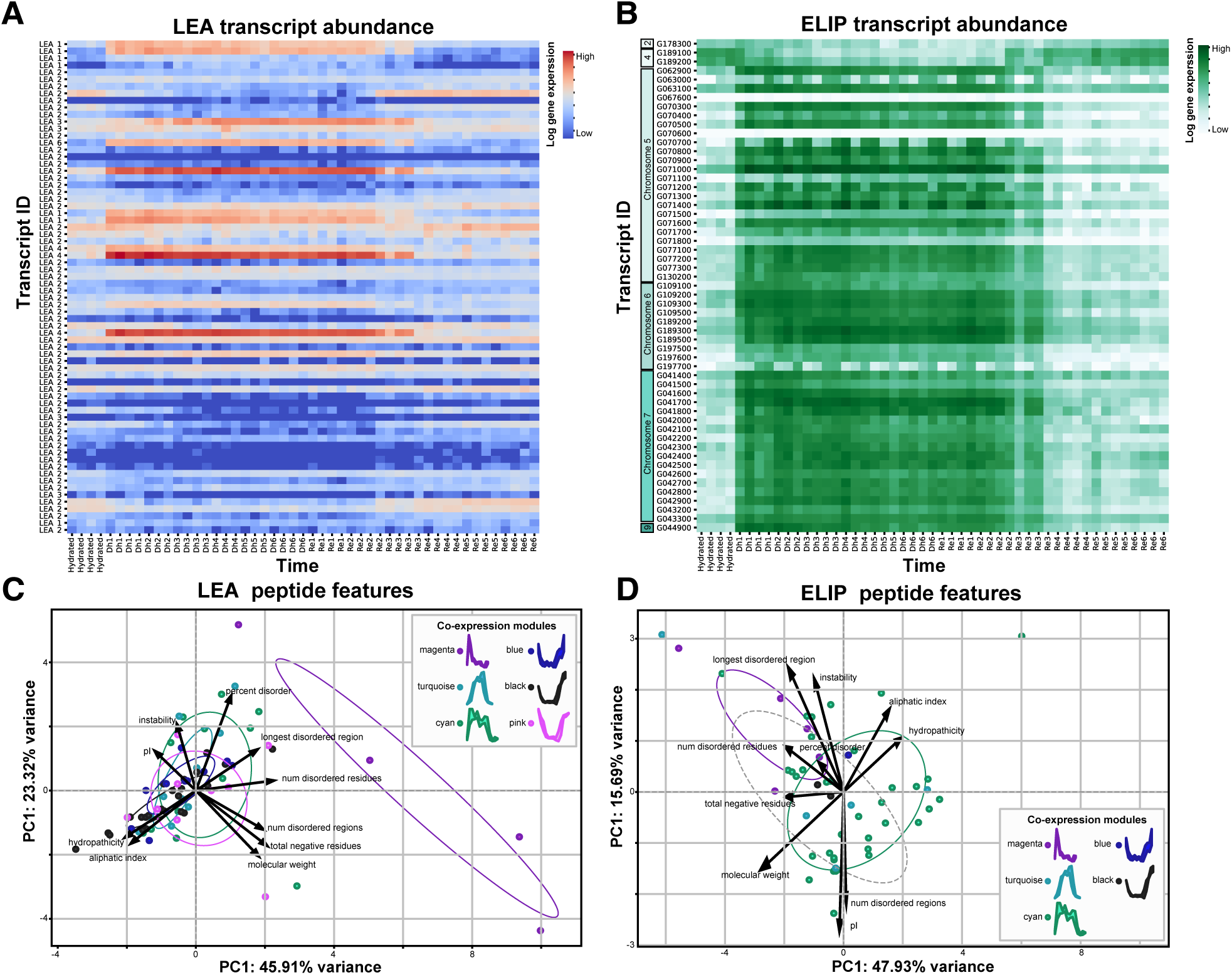
Expression dynamics of LEA and ELIP transcripts. Heatmaps of log transformed transcript abundance levels of (A) LEAs and (B) ELIPs across the time course. Data are only shown for plants that experienced full desiccation. Principal component analysis of peptide characteristics colored by module membership for (C) LEAs and (D) ELIPs.

To better understand the nature of these dynamic changes, we investigated differences in the disorder, hydropathicity, molecular weight, and charge of each predicted LEA and ELIP peptide and tested if any properties of the proteins were related to their expression profiles. In general, LEAs with higher hydropathic indices (more hydrophobic) were less disordered, suggesting a trade-off between hydrophobicity and flexibility (Figure 7C). More importantly, we found that the more disordered LEA proteins were expressed predominantly during desiccation (i.e., membership in magenta, cyan, turquoise coexpression modules), while less disordered LEAs exhibited reduced expression during this phase (i.e. blue, black, and pink modules). These trends were closely associated with LEA family classification, with LEA2 transcripts being less disordered and expressed at lower levels during dehydration compared to the highly disordered and more abundantly expressed LEA1 and LEA6 transcripts. We suspect that more disordered LEA transcripts may act as flexible molecular chaperones, stabilizing a broad array of cellular targets during desiccation, while less disordered variants may provide other functions. This functional and structural diversity within the LEA family points to their specialized roles in facilitating stress tolerance across different physiological contexts.

Our analyses revealed intriguing patterns within the ELIPs as well. Similar to other resurrection plants <u>(VanBuren et al. 2019)</u>, ELIP genes are expanded in tandem arrays in *M. flabellifolia* (Figure 7B). The majority of *M. flabellifolia* ELIPs exhibit the typical dehydration-induced increases in abundance (Figure 7B). However, a small number of these ELIPs with little to no expression under any condition and may be pseudogenes, and a few ELIPs are expressed only under hydrated conditions–especially those located in a small array on chromosome 4. Analysis of the peptide characteristics confirmed that ELIPs generally exhibit lower levels of intrinsic disorder compared to LEAs. Consequently, their expression patterns show weaker associations with peptide properties and most ELIPs were clustered in the cyan co-expression module, suggesting a coordinated role in desiccation response (Figure 7D). Interestingly, higher disorder in ELIPs appeared to be weakly linked with earlier induction during drying, hinting at a potential role in initiating protective mechanisms.

To better understand the drivers of differential expression of both ELIPs and LEAs we investigated the cis-regulatory elements (CREs) associated with each gene using the PLACE v30.0 database <u>(Higo et al. 1999)</u>. We tested for differences in CREs using between LEAs and ELIPs that were upregulated during dehydration (i.e., magenta, turquoise, and cyan modules) versus those that were downregulated during dehydration (i.e., blue, black, and pink modules). We identified multiple CREs that were significantly enriched in LEA genes upregulated during dehydration. The significant CREs included WBOXNTCHN48 (p = 0.0097), recognized by WRKY transcription factors that are involved in abiotic stress response in resurrection plants (Bakshi and Oelmüller, 2014; Xiang et al., 2021; Yang et al., 2023); CCA1ATLHCB1 (p = 0.0302), which is bound by the CCA1 transcription factor that interacts with light-harvesting complex genes (Wang et al., 1997; Wang and Tobin, 1998; Sun et al., 2019); IRO2OS (p = 0.0204) which is associated with the OsIRO2 transcription factor that regulates iron uptake and homeostasis (Ogo et al., 2006; Li et al., 2022); BOXIIPCCHS (p = 0.0273) which is a G-box-like motif essential for light regulation (Block et al., 1990); ACGTABREMOTIFA2OSEM (p = 0.0245) which is an ABA-responsive element (ABRE) associated with dehydration-induced gene expression (Narusaka et al., 2003; Zhang et al., 2024); and MYB1LEPR CRE (p = 0.0371) which is recognized by R2R3-MYB transcription factors and linked to ABA and drought-responsive gene regulation (Abe et al., 1997; Chakravarthy et al., 2003; Giarola et al., 2017).

There was only one CRE (CURECORECR; p = 0.0232) that was enriched in the LEA genes with downregulation during dehydration. This CRE is associated with metal and oxygen responses (Quinn and Merchant, 1995; Sharma et al., 2011) and is part of the G-box family of light-regulated elements, often recognized by bZIP and other transcription factors involved in photomorphogenesis and chloroplast development, suggesting that the LEAs not induced during dehydration may have roles in protecting against other abiotic stresses.

ELIPs showed weaker patterns of CRE enrichment. Only one CRE, TATCCACHVAL21 (*p* = 0.0443), was significantly enriched in the small set of ELIPs that were downregulated during dehydration. This element is a gibberellin-responsive motif linked to energy mobilization during germination (Gubler and Jacobsen, 1992; Isabel-LaMoneda et al., 2003; Martínez et al., 2005), suggesting that these ELIPs may retain canonical GA-linked functions in seed biology. Three additional CREs showed trends toward enrichment (*p* < 0.1), but did not pass the p-value threshold of < 0.05 required for significance. These included TATCCAYMOTIFOSRAMY3D (*p* = 0.0955), which was also more common in down-regulated ELIPs and is another GA-responsive element (Xie et al., 2016). Two CREs trended toward enrichment in the upregulated ELIPs: TAAAGSTKST1 (*p* = 0.0592), which is linked to guard cell function (Plesch et al., 2001), and MYCCONSENSUSAT (*p* = 0.0955), a MYC transcription factor site involved in ABA signaling and dehydration response (Abe et al., 2003), pointing towards ABA dependent stress induced expression.

## DISCUSSION

The chromosome-level, haplotype-resolved genome assembly of *M. flabellifolia* presented here provides a foundational resource for understanding the genomic architecture and resilience of this iconic resurrection plant. The *M. flabellifolia* genome exhibits strikingly uniform gene and repeat density across chromosomes and lacks clear evidence of conventional centromeres or pericentromeric regions. These features, along with a consistent 3D chromatin organization revealed by Hi-C, point toward a putative holocentric chromosome structure—an unusual and potentially adaptive genomic configuration that warrants further cytogenetic investigation. The two haplotypes are highly divergent, exhibiting substantial structural variation, including a large inversion on chromosome 1 and widespread gene presence-absence and copy number variation which may contribute to functional diversity in this dioecious outcrossing species.

We identified and characterized an XY sex determination system in *M. flabellifolia*, with heterogametic males and a small, ∼700 Kb SDR located on chromosome 8. The SDR contains just six gene models, including a kinesin-like gene previously implicated in anther development in rice (Zhou et al., 2011), along with two Y-specific genes of unknown function. This compact and minimally differentiated sex chromosome pair offers a rare opportunity to study the evolution of dioecy and sexual dimorphism in a non-model lineage under extreme environmental stress. Although no direct differences in desiccation tolerance between sexes have been reported, population-level studies reveal skewed sex ratios along environmental gradients, with males more common in arid environments and females more frequent in mesic ones (Marks et al., 2022).

We profiled a dozen plants during a natural dehydration-rehydration event in the field, making our study one of the first to survey an angiosperm resurrection plant directly in the field. In general, transcriptomic changes were tightly linked to changes in RWC, reflecting the central role of water availability in altering cellular processes. Early transcriptomic changes, such as the general suppression of photosynthesis and growth, parallel previous findings and point towards an energy-saving strategy that helps to minimize oxidative damage (Farrant, 2000; Pardo et al., 2020; Farrant and Hilhorst, 2021; Marks et al., 2023; VanBuren et al., 2023). As desiccation progressed, a transition to more specialized responses occurred, including the suppression of cell division and circadian rhythm, and a shift towards a more complete cessation of cellular activity and mechanisms to preserve molecular integrity. These findings generally align with the early and late stages of dehydration observed across desiccation-tolerant taxa, characterized by the early suppression of metabolic activity to conserve energy and minimize damage during desiccation (Farrant, 2000; Alejo-Jacuinde et al., 2020; Oliver et al., 2020). Co-expression analysis further highlighted the complexity of desiccation responses in *M. flabellifolia* and demonstrated an increasing response to drying as cellular dehydration increased. Modules enriched for strategies of maintaining cellular stability and energy production during drought such as carbohydrate metabolism, oxidative stress responses, and cell wall modifications were activated early during the dehydration process in all plants. These modules included many of the known hallmarks of desiccation tolerance, such as trehalose and raffinose biosynthesis, pointing towards the early activation of key protective strategies. However, modules with severity-specific activation, suggest that mechanisms of transcription and translation regulation are induced at later stages, and highlight an escalating response that is programmed to match the intensity of dehydration stress.

Changes in LEA and ELIP expression during dehydration and rehydration reinforce their central role in protecting cells under extreme dehydration (VanBuren et al., 2019; Kc et al., 2024), and point towards the importance of priming in stress responses and offer an important perspective on real-world adaptive responses in ecologically relevant contexts. Elevated expression of specific LEA and ELIP transcripts in hydrated plants suggests a baseline level of protection that may equip *M. flabellifolia* for rapid water loss that we were only able to detect due to our sampling in natural field conditions. During dehydration, expression shifted to increasingly disordered LEA and ELIP copies, pointing towards a transition of mechanisms and highlighting the subtle but important role of intrinsic disorder in providing protection.

The significant enrichment of WRKY-binding W-box motifs, ABA-responsive elements, and light-regulatory CREs in upregulated LEA genes suggests that dehydration-induced expression of these genes is mediated through well-established drought-responsive pathways (Abe et al., 1997; Xiang et al., 2021). The presence of ABA-responsive elements reinforces the role of an ABA-dependent transcriptional response, while the enrichment of G-box and CCA1-related elements suggests light may also play a role in modulating LEA gene expression under dehydration stress. Interestingly, only one significant CREs was identified in downregulated LEA genes, implying that dehydration responses in *M. flabellifolia* primarily involve induction rather than repression of LEA genes. This activation-based regulation aligns with the physiological role of LEA proteins, where their accumulation is thought to help stabilize cellular structures and prevent protein aggregation during desiccation (Kc et al., 2024).

CRE enrichments also point to functional divergence among ELIP genes in *M. flabellifolia*. The small subset of ELIP genes that were downregulated during dehydration appear to retain ancestral regulatory features (Hutin et al., 2003), including promoter motifs responsive to gibberellin and light, both of which are traditionally linked to seed germination and photomorphogenic development (Fleet and Sun, 2005). This pattern suggests that these downregulated ELIPs may fulfill a conserved role in early seedling establishment but are not required—or are actively repressed—during desiccation. In contrast, the majority of ELIPs are highly expressed during desiccation and lack enrichment for these classical GA motifs. Many of these upregulated ELIPs are found in tandem arrays on chromosomes 5, 6, and 7, consistent with a history of gene duplication and potential neofunctionalization. Several motifs associated with stress responses and protective functions trended toward enrichment, suggesting the evolution of desiccation-specific regulatory modules in these tandem arrays. Taken together, this supports a model in which ancestral ELIPs serve germination-related functions under GA and light control, while duplicated copies have acquired new regulatory sequences that allow them to function during desiccation.

Taken together this study showcases the unique genomics of *M. flabellifolia* and provides an in-depth characterization of the complex biology of desiccation tolerance. As one of only two extant genera in the early-diverging eudicot order Gunnerales, *Myrothamnus* occupies a key phylogenetic position for comparative evolutionary studies. The ability to survive complete desiccation, coupled with a dioecious reproductive system, and the production of antioxidant-rich secondary metabolites with known medicinal value, make *M. flabellifolia* a powerful system for linking evolutionary genomics, stress biology, and applied plant science. This genome not only enables deeper insight into the molecular mechanisms of desiccation tolerance and sex determination, but also provides a foundation for future research into the ecological strategies and biotechnological potential of resurrection plants.

## METHODS

### Study organism

*M. flabellifolia* is a dioecious resurrection plant in the early diverging eudicot lineage, Gunnerales. Myrothamnaceae contains only one genus with just two species in it; *M. flabellifolia* and *M. moschatus*. *M. moschatus* is endemic to Madagascar and *M. flabellifolia* is distributed throughout southern Africa in disjunct populations (Figure 1). Both species occupy a narrow ecological niche, restricted to rocky sites, with minimal soil, intense abiotic stresses (e.g., aridity, heat, irradiation), and low competition. *Myrothamnus sp.* are unique among resurrection plants as they are the only dioecious angiosperm resurrection plants, are large and woody growing up to ∼1.5M tall, and produce a robust profile of secondary compounds with important cultural history and medicinal applications.

### Field sites and sample collections

*M. flabellifolia* plants for the current study were collected from three distinct sites in southern Africa. These sites span substantial geographic distance and environmental variation, from extremely arid sites in Erongo, Namibia (-22.00922 S, 15.92595 E), to intermediate sites in Limpopo, South Africa (-23.7949, 28.0705), to mesic sites in Mpumalanga, South Africa (-25.30229 S, 30.50631 E). The locations of study sites were marked with a Garmin 64csx GPS, target plants were tagged, and leaf tissue was collected for downstream analyses. A single focal site in Limpopo, South Africa was selected for high-resolution transcriptomic profiling across a dehydration time course.

Initially, we sampled 10 plants across the three study sites. We collected leaf tissues from two male and two female plants each in Mpumalanga and Limpopo, South Africa. We collected leaf tissue from one male and one female from the site in Namibia. Dry tissue from each plant was shipped to the USA under permit number P37-19-01892 and according to the MTA established between Drs. Farrant, VanBuren, and Marks. Leaf tissues were rehydrated, harvested, and flash frozen in liquid nitrogen. Genomic DNA was extracted and processed for Illimina DNAseq as described in detail below. The resulting sequence data were analyzed to estimate diversity across populations, heterozygosity, and to identify the sex determination region of *M. flabellifolia* (also described in detail below).

Based on these analyses, we targeted a single male plant from Limpopo South Africa for reference genome sequencing and assembly. We selected a male plant because our analysis indicated that *M. flabellifolia* has an XY sex determination system and sequencing the heterogametic sex would thus provide a more comprehensive assessment of the genome sequence. For the reference genome assembly, we collected additional healthy green tissue from the reference accession (var. SSDT_37) in the field and immediately flash-froze it in liquid nitrogen. Frozen leaf tissue was shipped to the USA under USDA permit number P37-19-01892 and according to the MTA established between Drs. Farrant, VanBurnen, and Marks.

### DNA extraction and library preparation

High molecular weight (HMW) DNA was extracted from the 10 accessions and the reference genotype (var. SSDT_37) following standard protocols. In short, high molecular weight DNA was extracted using the protocol of <u>Doyle and Doyle (1987)</u> with minor modifications. Flash-frozen biomass was ground to a fine powder in a frozen mortar with liquid nitrogen followed by very gentle extraction in 2% CTAB buffer (that included proteinase K, PVP-40 and beta-mercaptoethanol) for 30 min to 1h at 50 °C. After centrifugation, the supernatant was transferred to a new tube, treated with 200ul 50mM PSMF for 10 minutes at room temperature then gently extracted twice with 24:1 Chloroform: Isoamyl alcohol. The upper phase was transferred to a new tube and 1/10th volume 3 M Sodium acetate was added, gently mixed, and DNA precipitated with iso-propanol. DNA precipitate was collected by centrifugation, washed with 70% ethanol, air dried for 5-10 minutes and dissolved thoroughly in an elution buffer at room temperature followed by RNAse treatment. DNA purity was measured with Nanodrop, DNA concentration measured with Qubit HS kit (Invitrogen) and DNA size was validated by Femto Pulse System (Agilent).

Illumina sequencing libraries were prepared by shearing 500 ng - 1.5 ug HMW DNA on a Covaris instrument to 350 bp. DNA fragments were bead cleaned, end repaired, and size selected to remove large and small fragments. After adenylation, adaptors were ligated according to the Illumina TruSeq PCR-Free DNA Library Prep Kit. Libraries were sequenced on an Illumina 6000 Instrument PE150 (see Supplementary Table S1). PacBio DNA sequencing libraries were prepared by shearing HMW DNA using a megaruptor shearing device (Diagenode), ligating SMRTbell adaptors (SMRTbell Prep Kit v3.0) and sizing the final library using a BluePippin Instrument (SAGE Science) at 10-50kb. Omni-C libraries were prepared by grinding frozen leaf tissue using a freezer mill followed by library construction using a Dovetail-C kit (Cantata Bio) (see Supplementary Table S2).

### Illumina and PacBio genome Sequencing

We sequenced the genome of *M.flabellifolia* using a whole genome shotgun sequencing strategy according to standard sequencing protocols. Sequencing reads were generated using Illumina and PACBIO platforms. Illumina and PACBIO reads were sequenced at the HudsonAlpha Genome Sequencing Center in Huntsville, Alabama. Illumina reads were sequenced using the Illumina NovoSeq6000 platform. PacBio Sequencing primer was annealed to the SMRTbell template library and sequencing polymerase was bound to them using Sequel II Binding kit 2.0. The prepared SMRTbell template library was then sequenced on a Pacific Biosciences Sequel IIe sequencer using 8M v1 SMRT cells and Version 2.0 sequencing chemistry with 1×1800 sequencing movie run times. Two 400bp insert 2×150 Illumina fragment libraries were sequenced with a coverage of 89.28X along with one 2×150 HiC library with a coverage of 56.69x (see Supplementary Table S1) sequenced on a NovaSeq 6000 Instrument PE150. Prior to assembly, Illumina fragment reads were screened for phix contamination.

Reads composed of > 95% simple sequence were removed. Illumina reads < 50bp after trimming for adapter and quality (q < 20) were removed. The final read set consisted of 1,592,906,904 reads for a total of 89.28X coverage of high-quality Illumina bases. For the PACBIO sequencing, a total raw sequence yield of 110.49 Gb, with a total coverage of 42.99X per haplotype, was generated (see Supplementary Table S2).

### Genome assembly and construction of pseudomolecule chromosomes

HAP1 and HAP2 version 1.0 assemblies were generated by assembling the 6,672,299 PACBIO CCS reads (42.99X per haplotype) using the HiFiAsm+HIC assembler (Cheng et al., 2021) and subsequently polished using RACON (Vaser et al., 2017). This produced initial assemblies of both haplotypes. The HAP1 assembly consisted of 370 scaffolds (370 contigs), with a contig N50 of 13.6 Mb, and a total genome size of 1,290.4 Mb (see Supplementary Table S3). The HAP2 assembly consisted of 333 scaffolds (333 contigs), with a contig N50 of 10.9 Mb, and a total genome size of 1,295.2 Mb (see Supplementary Table S4).

To improve these initial assemblies, Hi-C Illumina reads from *M. flabellifolia* reference accession (var. SSDT_37), were separately aligned to the HAP1 and HAP2 contig sets with Juicer (Durand et al., 2016), and chromosome scale scaffolding was performed with 3D-DNA (Dudchenko et al., 2017). No misjoins were identified in either the HAP1 or HAP2 assemblies. The contigs were then oriented, ordered, and joined together into 10 chromosomes per haplotype using the HiC data. A total of 154 joins were applied to the HAP1 assembly, and 179 joins for the HAP2 assembly. Each chromosome join was padded with 10,000 Ns. Contigs terminating in significant telomeric sequence were identified using the (TTTAGGG)_n_ repeat, and care was taken to make sure that they were properly oriented in the production assembly. The remaining scaffolds were screened against bacterial proteins, organelle sequences, GenBank nr, and removed if found to be a contaminant. After forming the chromosomes, it was observed that some small (<20Kb) redundant sequences were present on adjacent contig ends within chromosomes. To resolve this issue, adjacent contig ends were aligned to one another using BLAT <u>(Kent 2002)</u>, and duplicate sequences were collapsed to close the gap between them. A total of 9 adjacent contig pairs were collapsed in the HAP1 assembly and 3 in the HAP2 assembly. The *M. flabellifolia* chloroplast and mitochondrial genomes were assembled with the OatK pipeline (https://github.com/c-zhou/oatk) utilizing the angiosperm database. The chloroplast and mitochondrial genomes were then polished using RACON (Vaser et al., 2017) and included as part of the release.

Finally, homozygous SNPs and INDELs were corrected in the HAP1 and HAP2 releases using ∼54X of Illumina reads (2×150, 400bp insert) by aligning the reads using BWA-MEM (Li, 2013) and identifying homozygous SNPs and INDELs with the GATK’s UnifiedGenotyper tool (McKenna et al., 2010). A total of 920 homozygous SNPs and 9,867 homozygous INDELs were corrected in the HAP1 release, while a total of 862 homozygous SNPs and 10,258 homozygous INDELs were corrected in the HAP2 release. The final version 1.0 HAP1 release contained 1,285.3 Mb of sequence, consisting of 195 contigs with a contig N50 of 11.4 Mb and a total of 99.99% of assembled bases in chromosomes (see Supplementary Table S5). The final version 1.0 HAP2 release contains 1,274.7 Mb of sequence, consisting of 169 contigs with a contig N50 of 13.7 Mb and a total of 100% of assembled bases in chromosomes (see Supplementary Table S6).

Completeness of the euchromatic portion of the version 1.0 assemblies was assessed using an rnaSEQ library (library ID: HOSHY). The aim of this analysis is to obtain a measure of completeness of the assembly, rather than a comprehensive examination of gene space. The transcripts were aligned to the assembly using BWA-MEM (Li, 2013). The screened alignments indicate that 98.89% of the rnaSEQ reads aligned to the HAP1 version 1.0 release, and 98.89% aligned to the HAP2 version 1.0 release.

### Genome annotation

Transcript assemblies were made from ∼455M pairs of 2×150 stranded paired-end Illumina RNA-seq reads using PERTRAN, which conducts genome-guided transcriptome short read assembly via GSNAP (Wu and Nacu, 2010) and builds splice alignment graphs after alignment validation, realignment, and correction. To obtain ∼663,000 and ∼664,000 (for HAP1 and HAP2, respectively) putative full-length transcripts, about 15M PacBio Iso-Seq CCSs were corrected and collapsed by a genome-guided correction pipeline. The pipeline aligns CCS reads to the genome with GMAP (Wu and Nacu, 2010), corrects small indels in splice junctions, and clusters alignments when all introns are the same or ≥ 95% overlap for single-exon alignments.

Subsequently, PASA (Haas et al., 2003) was used to construct 519,584 and 517,780 (for HAP1 and HAP2, respectively) transcript assemblies by combining the transcript assembly sets described above.

A repeat library was created from *de novo* repeats predicted by RepeatModeler2 (Flynn et al., 2020) on the *M. flabellifolia* var. *SSDT_37* HAP1 v1.0 genome. The predicted repeats underwent functional analysis through InterProScan (Jones et al., 2014), incorporating the Pfam (Mistry et al., 2021) and PANTHER (Mi et al., 2019) databases. Any repeats that displayed significant hits to protein-coding domains were subsequently excluded from the repeat set.

Finally, the constructed species-specific repeat library was used to soft-mask both haplotypes with RepeatMasker (Flynn et al., 2020).

Putative gene loci were determined by transcript assembly alignments and/or EXONERATE (Slater and Birney, 2005) alignments of proteins from *Mimulus guttatus*, *Lactuca sativa*, *Hydrangea quercifolia*, *Kalanchoe fedtschenkoi*, *Arabidopsis thaliana*, *Vitis vinifera*, *Gossypium hirsutum*, *Populus trichocarpa*, *Glycine max*, *Prunus persica*, *Liriodendron tulipifera*, *Eschscholzia californica*, *Solanum lycopersicum*, *Beta vulgaris*, *Brassica rapa*, *Citrus sinensis*, *Medicago truncatula*, *Sorghum bicolor*, *Oryza sativa*, *Osyris compressa*, *Macadamia integrifolia*, *Telopea speciosissima*, *Nelumbo nucifera*, and Swiss-Prot release 2022_04 of eukaryotic proteomes to repeat-soft-masked *M. flabellifolia* var. *SSDT_37* v1.0 HAP1 and HAP2 genomes with up to 2,000 BP extension on both ends unless extending into another locus on the same strand. Gene models in each locus were predicted by homology-based predictors, FGENESH+ <u>(Salamov and Solovyev 2000)</u>, FGENESH_EST (similar to FGENESH+, but using EST to compute splice site and intron input instead of protein/translated ORF), EXONERATE, PASA assembly ORFs (in-house homology constrained ORF finder), and AUGUSTUS (Stanke et al., 2006) trained on the high confidence PASA assembly ORFs and with intron hints from short read alignments. The best-scored predictions for each locus were selected using multiple positive factors including EST and protein support, and one negative factor: overlap with repeats. The selected gene predictions were improved by PASA. The improvement included adding UTRs, splicing correction, and adding alternative transcripts.

PASA-improved gene model proteins were subjected to protein homology analysis to the above-mentioned proteomes to obtain a Cscore and protein coverage. Cscore is a protein BLASTP score ratio to the mutual best hit BLASTP score and protein coverage is the percentage of protein aligned to the best of homologs. PASA-improved transcripts were selected based on Cscore, protein coverage, EST coverage, and their CDS overlap with repeats. The transcripts were selected if their Cscore and protein coverage were ≥ 0.5 or if covered by ESTs. For gene models whose CDS overlapped repeats by more than 20%, their Cscore had to be at least 0.9 and homology coverage at least 70% to be selected. The selected gene models were subject to Pfam analysis and gene models without strong transcriptome and homology support whose proteins were more than 30% overlapped by Pfam TE domains were removed. Incomplete gene models, low homology supported without fully transcriptome-supported gene models, short single exon (< 300 BP CDS) without protein domains nor good expression, and repetitive gene models without strong homology support were manually filtered out.

TRASH was used to annotate repetitive sequences for the identification of putative centromeric regions in *M. flabellifolia*. Ultimately, we did not observe distinct clustering of satellite sequences and instead found satellite repeats to be distributed across the genome.

Genomes used in the analysis were fetched from phytozome *M. flabellifolia* var. SSDT_37 HAP1 and HAP2. The most common repeat annotated by TRASH was a 7 bp satellite that occurs regularly and is wide-spread across the genome. Given that this 7bp satellite is so wide-spread, we additionally checked for any structure of the next most frequent satellites identified by TRASH. For the next 8 most frequent satellites identified by TRASH (26 bp, 34 bp, 35, bp 49 bp, 63 bp, 64 bp, 65 bp, and 66 bp), we did not observe any obvious signals of clustering and the distribution of satellite sequences were spread around the genome. Four satellite sequences are plot per graphic for increased visibility of their distribution.

### Sex determination region

We used resequence data from 10 *M. flabellioflia* accessions (5 females, 5 males) to identify kmers unique to males (Y-mers; (Carey et al., 2024a, b)). We filtered the data using Trimmomatic v0.39 with leading and trailing values of 3, sliding window of 30, jump of 10, and a minimum remaining read length of 40 (Bolger et al., 2014a) and identified k-mers using meryl v1.3 *count* (Rhie et al., 2020). We used meryl’s *intersect* function to identify all 21-mers shared in females, followed by *difference* to identify the 21-mers only found in males. Y-mers were mapped to the assemblies in order to identify putatively Y-associated contigs using BWA-MEM v0.7.17 (Li and Durbin, 2009; Li, 2013) with flags ‘-k 21’, ‘-T 21’, ‘-a’, and ‘-c 10’. To determine which genes were X-or Y-specific, we used OrthoFinder v2.5.2 (Emms and Kelly, 2015, 2019)), to identify homologs.

### Field sampling and physiological measurements across desiccation time course

We tracked 12 plants from the focal population in Limpopo South Africa across a natural dehydration-rehydration event in the field. Initially, study plants were tagged and phenotyped to quantify plant height, sex, and flower number. These plants were then monitored throughout a natural dehydration and rehydration event spanning 6 days in the field. Each plant was sampled at baseline hydrated conditions (14h00 on Jan 14th, 2020) and at 6 progressive dehydration timepoints. Dehydration sampling was done at 44 hours (10h00, Jan 16th, 2020), 48 hours (14h00, Jan 16th, 2020), 52 hours (18h00, Jan 16th, 2020), 68 hours (10h00, Jan 17th, 2020), 72 hours (14h00, Jan 17th, 2020) and 76 hours (18h00, Jan 17th, 2020) after sampling began. We then waited for a natural rainfall event and sampled rehydration timepoints at 2 hours (9h00, Jan 18th, 2020), 4 hours (11h00, Jan 18th, 2020), 8 hours (15h00, Jan 18th, 2020), 12 hours (19h00, Jan 18th, 2020), 24 hours (9h00, Jan 19th, 2020), and 48 hours (9h00, Jan 20th, 2020) after rainfall. Environmental data on temperature, relative humidity, and rainfall were tracked with a Hobo weather station and U23 pro V2 dataloggers. At each sampling timepoint, we quantified the RWC of the sample and flash froze leaf tissue in liquid nitrogen for downstream RNA extractions. RWC was quantified by measuring the mass of 10-15 leaves from each plant. Leaf mass was weighed immediately after collection (fresh weight), again after 48 hours submerged in *d*H20 in darkness at 4°C (turgid weight), and finally after 48 hours in a 70°C drying oven (dry weight). RWC was calculated as (fresh weight - dry weight)/(turgid weight - dry weight). Over the course of the experiment, individual plants dried at different rates and we assigned them to groups based on how much they dried during the time course. Plants were assigned to either “mild dehydration”, “severe dehydration”, or “desiccation” groups according to their minimum RWC during the time course.

### RNA extraction and sequencing

RNA was extracted from a total of 149 samples from 12 individuals across 13 timepoints. Three additional samples from flower tissue were included to aid in genome annotation. RNA was extracted using the Spectrum total plant RNA kit according to manufacturer’s instructions, with on-column DNAse treatment included in the pipeline. RNA samples were further processed to remove impurities and contaminants using the Zymo RNA clean and concentrate kit according to manufacturer’s instructions.

Plate-based RNA library prep was performed on the PerkinElmer Sciclone NGS robotic liquid handling system using Illumina’s TruSeq Stranded mRNA HT sample prep kit utilizing poly-A selection of mRNA following the protocol outlined by Illumina in their user guide (https://support.illumina.com/sequencing/sequencing_kits/truseq-stranded-mrna.html). Briefly, total RNA starting material was 1000 ng per sample and 8 cycles of PCR were used for library amplification. The prepared libraries were quantified using KAPA Biosystems’ next-generation sequencing library qPCR kit and run on a Roche LightCycler 480 real-time PCR instrument.

Sequencing of the flowcell was performed on the Illumina NovaSeq sequencer using NovaSeq XP V1.5 reagent kits, S4 flowcell, following a 2×151 indexed run recipe. RNA libraries were sequenced on an Illumina NovaSeq S4 for 150bp PE reads.

Long-read RNAseq (isoSeq) data were generated for 12 leaf and flower samples from the reference accession (var. SSDT_37) to aid in genome annotation. Full-length cDNA was synthesized using template switching technology with NEBNext Single Cell/Low Input cDNA Synthesis & Amplification Module kit. The first-strand cDNA was amplified and multiplexed with NEBNext High-Fidelity 2X PCR Master Mix using Barcoded cDNA PCR primers. The amplified cDNA was purified using 1.3X ProNex beads for non-size selection or 0.89X ProNex beads for above 2 kb size selection and liked sizes were pooled at the equimolar ratios in a designated degree-of-pool in the worksheet using PacBio Multiplexing Calculator. The pooled samples were end-repaired, A-tailed and ligated with overhang non-barcoded adaptors using SMRTbell Express 2.0 kit. IsoSeq libraries were sequenced on a PacBioSequel II.

### Transcriptomic analyses

The resulting RNAseq reads were processed to quantify transcript abundance following a pipeline developed by the VanBuren Lab (https://github.com/pardojer23/RNAseqV2). Briefly, read quality was assessed with fastQC (v 0.23) and reads were trimmed with trimmomatic (v 0.38) (Bolger et al., 2014b) to remove adapters and low quality bases. Trimmed reads were sudo-aligned to the reference genome using Salmon (v 1.9.0) (Patro et al., 2017), and the resulting quantification files were processed with tximport (v 3.18) (Soneson et al., 2015) to generate raw count and transcripts per million (TPM) expression matrices. Hierarchical clustering was conducted for basic quality control and visualization of sample relationships across experimental timepoints and biological replicates. A principal component analysis (PCA) of TPM values was used to further visualize replicate and sample relationships.

While PCA provided some level of dimensionality reduction, residual heterogeneity, experimental differences, noise, or genotype level differences in the dataset might have obscured underlying biology. To address this, we applied topological data analysis (TDA) using the Mapper algorithm <u>(van Veen et al. 2019; Palande et al. 2023; Marks et al. 2024)</u>, which provides a flexible and scalable approach for exploring high-dimensional, sparse datasets.

Mapper requires a lens function, a user-defined feature that shapes how the data are clustered and connected. We used RWC as the lens, anchoring it to the fully hydrated condition, to reflect physiological water status across samples. This allowed us to explore expression changes along a continuum of dehydration and recovery. For our mapper graph, we specified 110 intervals with a 90% overlap. The topology was plotted to visualize relationships across samples.

Differentially abundant transcripts (DATs) were identified with the DEseq2 R package (v 1.42.0) (Love et al., 2014). Initially, we tested multiple models for identifying DATs in DEseq2, including a model that identified DATs by pairwise comparisons of each timepoint against well-watered and a model that used the continuous variable of RWC as a covariate. DATs identified by pairwise comparisons were summarized into a nonredundant list of up and down regulated genes during dehydration and rehydration. To select the best performing model, we quantified similarities and differences in the number and identity of DATs defined by each model using Venn diagrams. There was a high degree of overlap in DATs identified by both models.

Ultimately, we selected the model based on pairwise comparisons for downstream analyses because it provided differentiation between the effects of time and drying rate, whereas the model based on RWC did not account for the effect of time.

We identified DATs separately for plants in each of the three drying groups (fast, intermediate, and slow) and compared the overlap in DATs between groups using Venn diagrams. We then investigated the functional significance of the shared and unique DATs using gene ontology (GO) enrichment analysis. TopGO (Rahnenfuhrer, 2019) was used to identify enriched GO terms within sets of unique and shared DATs. We further visualized changes in DATs across the time course with Alluvial diagrams generated with the alluvial R package (v 0.1-2) (Edwards and Bojanowski, 2016).

While DAT analyses can be informative for identifying and describing overarching patterns and large shifts in transcript abundance, more nuanced patterns of gene expression and regulation can be obscured in classical differential analysis. To gain insight into these aspects, we generated co-expression networks using Weighted Gene Co-expression Network Analysis (WGCNA) R package (v1.7) (Langfelder and Horvath, 2008). Co-expression modules were identified using the complete transcript abundance matrix of all genes in all samples. To construct the co-expression network, we first determined a soft thresholding power for the dataset. The thresholding power was chosen to satisfy WGCNA’s assumption that a weighted co-expression network is scale-free. An adjacency matrix, representing the strength of connections between genes in the network, was constructed using the selected soft thresholding power. This matrix was then converted to a topological overlap matrix (TOM) and hierarchal clustering was used on the TOM to group genes into modules based on similar expression patterns. We plotted the expression profiles of each module for each of the drying groups separately. This allowed us to visualize differences between the drying groups for each gene expression module. We then investigated the functional significance of modules using GO enrichment analysis, implemented with the R package TopGO (Rahnenfuhrer, 2019).

Both Late Embryogenesis Abundant (LEA) and Early Light Induced Protein (ELIP) gene families have important roles in desiccation tolerance (VanBuren et al., 2019; Hernández-Sánchez et al., 2022), and we conducted targeted analyses to quantify the expression dynamics of these gene families. LEA proteins were identified by the genome annotation pipeline described above, and ELIPs were identified via a BLAST search against the Arabidopsis ELIP1 gene. We plotted the normalized expression of each LEA and ELIP paralogs in each sample to visualize expression patterns across the time course in heatmaps. The predicted peptide sequences of each LEA and ELIP transcript were analysed on ExPASy <u>(Gasteiger et al. 2003)</u> to retrieve characterisation information and on pondr <u>(Xue et al. 2010)</u> for disorder prediction. ClustalOmega <u>(Sievers et al. 2011)</u> used to generate multiple sequence alignments of sequences and to retrieve percent identity per protein group using default settings.

To investigate the cis-regulatory landscape of LEA gene expression during dehydration in *M. flabellifolia*, we first extracted the 1 kb upstream promoter regions of all LEA genes identified in co-expression modules. Promoter coordinates were determined using the annotated GFF3 file for the *M. flabellifolia* (var. SSDT_37) genome, and strand orientation was taken into account to extract the appropriate upstream regions using custom `awk` commands. Promoter sequences were retrieved from the reference genome using BEDtools v2.30.0 (Quinlan and Hall, 2010).

Cis-regulatory elements (CREs) within these promoter regions were then identified using the PLACE v30.0 database <u>(Higo et al. 1999)</u>. Each CRE’s presence or absence was recorded as a binary matrix, where rows represented LEA genes and columns represented individual CREs.

LEA genes were grouped based on their co-expression module assignment: turquoise, magenta, and cyan modules were categorized as upregulated during dehydration; while black, blue, and pink modules were categorized as downregulated.

To determine whether specific CREs were enriched in the promoters of upregulated or downregulated LEA genes, we performed logistic regression analysis using R. Each CRE was tested individually for its association with gene expression group (upregulated vs. downregulated) using a binomial generalized linear model. The dependent variable was module group (downregulated vs upregulated), and the independent variable was CRE presence vs absence. Odds ratios and p-values were extracted for each CRE. Odds ratios greater than 1 indicate enrichment in upregulated genes, while odds ratios less than 1 indicate enrichment in downregulated genes. CREs with p-values < 0.05 were considered statistically significant. We used this information to identify regulatory motifs that may contribute to LEA transcript abundance patterns in response to dehydration.

## AUTHOR CONTRIBUTIONS

RAM, JMF, RV, and JLM conceived of the study. RAM, JT, KB, and JG collected and generated the data. RAM, SBC, JTL, AH, TB, NMC, JS, AL, CP, JY, DB, JW and JWJ analyzed the data. RAM, SBC, JTL, LVDP, NC, JB, AH, RV, JLM, and JMF interpreted the data. RAM wrote the manuscript with sections contributed by SBC, JTL, TB, JB, and LVDP. All authors reviewed and approved the final manuscript.

## DATA AVAILABILITY

Genome assemblies and annotations are available at NCBI under bioproject PRJNA1114142 and at the Department of Energy’s Joint Genome Institute genome portal: https://phytozome.jgi.doe.gov/info/Mflabellifoliavar_SSDT_37HAP1_v1_1 and https://phytozome.jgi.doe.gov/info/Mflabellifoliavar_SSDT_37HAP2_v1_1 as part of the open green genomes project (https://phytozome.jgi.doe.gov/ogg). DNA sequencing reads are deposited in NCBI’s SRA under the umbrella project BioProjectPRJNA1114756. All RNA sequencing reads have been deposited in NCBI’s SRA under BioProjects PRJNA1114143-PRJNA1114253, and accession numbers SRR29328105, SRR29328106, SRR29328107, SRR29328108, SRR29328109, SRR29328110, SRR29328111, SRR29328112, SRR29328113, SRR29328114, SRR29328115, SRR29328116, SRR29328117, SRR29328119, SRR29328120, SRR29328121, SRR29328122, SRR29328123, SRR29328124, SRR29328125, SRR29328126, SRR29328127, SRR29328128, SRR29328129, SRR29328130, SRR29328131, SRR29328132, SRR29328133, SRR29328134, SRR29328135, SRR29328136, SRR29328137, SRR29328138, SRR29328139, SRR29328140, SRR29328141, SRR29328142, SRR29328143, SRR29328144, SRR29328145, SRR29328170, SRR29328171, SRR29328172, SRR29328173, SRR29328174, SRR29328175, SRR29328176, SRR29328177, SRR29328178, SRR29328179, SRR29328180, SRR29328181, SRR29328182, SRR29328183, SRR29328199, SRR29328200, SRR29328201, SRR29328202, SRR29328203, SRR29328204, SRR29328211, SRR29328212, SRR29328213, SRR29328214, SRR29328220, SRR29328222, SRR29328223, SRR29377105, SRR29377106, SRR29377107, SRR29377108, SRR29377109, SRR29377110, SRR29377111, SRR29377112, SRR29377113, SRR29377114, SRR29377115, SRR29377116, SRR29377117.

## ACKNOWLEDGEMENTS

This work was funded by NSF IOS-PRFB-1906094 to RAM, DBI-2213983 to RAM and RV, MCB-1817347 to RV, and IOS-PGRP CAREER-223930 to AH. JMF acknowledges the South African DSI and NRF, grant number 98406. NC thanks the NRF for her PhD scholarship. We thank landowners and stewards Ken Maude, Pieter and Nadine Vervoort, John and Sandy Burrows, and Jennie and Pieter Pretorious for permission to collect plants, and Wayne and Messiah Mudenda for assistance with field logistics. We thank the University of Cape Town for access to facilities and Keren Cooper for logistical support. We also thank the South African National Herbarium, Pretoria for assistance identifying and vouchering of specimens and the United States Department of Agriculture for providing import permits. The work (proposal: 10.46936/10.25585/60001405) conducted by the U.S. Department of Energy Joint Genome Institute (https://ror.org/04xm1d337), a DOE Office of Science User Facility, is supported by the Office of Science of the U.S. Department of Energy operated under Contract No. DE-AC02-05CH11231.

## Notes

### Competing Interest Statement

The authors have declared no competing interest.

### Summary of Updates

the figures in the original submission appeared in the wrong order.

