## SUPPLEMENTARY MATERIAL for "The architecture of resilience: a genome assembly of *Myrothamnus flabellifolia* sheds light on desiccation tolerance and sex determination"

**Construction of the scaffold assembly.** A total of 6,672,299 PACBIO reads (42.99x per haplotype) were assembled using HiFiAsm+HIC assembler (Cheng et al., 2021), and formed the starting point of the version 1.0 release. The 1,592,906,904 Illumina fragment 2x150 reads (89.28x sequence coverage) was used for fixing homozygous snp/indel errors in the consensus. Chromosomes were scaffolded using the 1,047,438,958 2x150 (56.69x) HiC reads.

**Table S1.** Genomic libraries included in the *Myrothamnus flabellifolia (var. SSDT_37)* genome assembly and their respective assembled sequence coverage levels in the final release.

^*^Average read length of PACBIO reads.

| **Library** | **Sequencing Platform** | **Average Read/Insert Size** | **Read Number** | **Assembled Sequence Coverage (x)** |
| --- | --- | --- | --- | --- |
| JCHZ_1024 | Illumina (2x150) | 400 | 866,865,500 | 48.46 |
| JCHZ_1027 | Illumina (2x150) | 400 | 726,041,404 | 40.82 |
| KIQF | Illumina-HiC  (2x150) | N/A | 1,047,438,958 | 56.69 |
| HSZWT | PACBIO | 15,920^*^ | 6,672,299 | 42.99 |
| **Total** |  | N/A | 2,647,018,161 | 188.96 |

**Table S2.** PACBIO CCS library statistics for the libraries included in the *Myrothamnus flabellifolia (var. SSDT_37)* genome assembly and their respective assembled sequence coverage levels.

| **Cutoff** | **Number of Reads** | **Basepairs** | **Average Read Length** | **Coverage** |
| --- | --- | --- | --- | --- |
| 0 | 6,672,299 | 110,486,722,577 | 15,920 | 42.99x |
| 1,000 | 6,668,201 | 110,484,391,751 | 15,923 | 42.99x |
| 2,000 | 6,662,596 | 110,475,853,380 | 15,927 | 42.99x |
| 3,000 | 6,655,523 | 110,458,025,667 | 15,932 | 42.98x |
| 4,000 | 6,645,888 | 110,424,127,530 | 15,939 | 42.97x |
| 5,000 | 6,634,158 | 110,371,211,854 | 15,948 | 42.94x |
| 6,000 | 6,620,629 | 110,296,607,955 | 15,958 | 42.92x |
| 7,000 | 6,606,535 | 110,205,086,898 | 15,969 | 42.88x |
| 8,000 | 6,592,903 | 110,102,827,301 | 15,979 | 42.84x |
| 9,000 | 6,577,630 | 109,972,808,781 | 15,990 | 42.79x |
| 10,000 | 6,555,731 | 109,763,856,821 | 16,006 | 42.71x |
| 11,000 | 6,501,178 | 109,184,346,854 | 16,047 | 42.48x |
| 12,000 | 6,179,853 | 105,452,613,236 | 16,290 | 41.03x |
| 13,000 | 5,512,543 | 97,094,640,388 | 16,822 | 37.78x |
| 14,000 | 4,747,524 | 86,764,891,354 | 17,486 | 33.76x |
| 15,000 | 3,988,602 | 75,763,555,955 | 18,225 | 29.48x |
| 16,000 | 3,282,195 | 64,819,629,728 | 19,011 | 25.22x |
| 17,000 | 2,650,337 | 54,401,409,829 | 19,828 | 21.17x |
| 18,000 | 2,104,845 | 44,862,857,820 | 20,660 | 17.46x |
| 19,000 | 1,645,614 | 36,374,240,625 | 21,496 | 14.15x |

**Table S3**. Summary statistics of the initial output of the HAP1 RACON polished HiFiAsm+HIC assembly. The table shows total contigs and total assembled basepairs for each set of scaffolds greater than the size listed in the left hand column.

| **Minimum**  **Scaffold**  **Length** | **Number of**  **Scaffolds** | **Number of**  **Contigs** | **Scaffold Size** | **Basepairs** | **% Non-gap Basepairs** |
| --- | --- | --- | --- | --- | --- |
| 5 Mb | 84 | 84 | 1,097,947,157 | 1,097,947,157 | 100.00% |
| 2.5 Mb | 116 | 116 | 1,217,549,339 | 1,217,549,339 | 100.00% |
| 1 Mb | 144 | 144 | 1,265,106,648 | 1,265,106,648 | 100.00% |
| 500 Kb | 154 | 154 | 1,272,868,595 | 1,272,868,595 | 100.00% |
| 250 Kb | 160 | 160 | 1,275,059,470 | 1,275,059,470 | 100.00% |
| 100 Kb | 184 | 184 | 1,278,798,069 | 1,278,798,069 | 100.00% |
| 50 Kb | 370 | 370 | 1,290,360,047 | 1,290,360,047 | 100.00% |
| 25 Kb | 370 | 370 | 1,290,360,047 | 1,290,360,047 | 100.00% |
| 10 Kb | 370 | 370 | 1,290,360,047 | 1,290,360,047 | 100.00% |
| 5 Kb | 370 | 370 | 1,290,360,047 | 1,290,360,047 | 100.00% |
| 2.5 Kb | 370 | 370 | 1,290,360,047 | 1,290,360,047 | 100.00% |
| 1 Kb | 370 | 370 | 1,290,360,047 | 1,290,360,047 | 100.00% |
| 0 bp | 370 | 370 | 1,290,360,047 | 1,290,360,047 | 100.00% |

**Table S4**. Summary statistics of the initial output of the HAP2 RACON polished HiFiAsm+HIC assembly. The table shows total contigs and total assembled basepairs for each set of scaffolds greater than the size listed in the left hand column.

| **Minimum**  **Scaffold**  **Length** | **Number of**  **Scaffolds** | **Number of**  **Contigs** | **Scaffold Size** | **Basepairs** | **% Non-gap Basepairs** |
| --- | --- | --- | --- | --- | --- |
| 5 Mb | 91 | 91 | 1,067,419,368 | 1,067,419,368 | 100.00% |
| 2.5 Mb | 130 | 130 | 1,214,090,884 | 1,214,090,884 | 100.00% |
| 1 Mb | 163 | 163 | 1,269,161,900 | 1,269,161,900 | 100.00% |
| 500 Kb | 179 | 179 | 1,280,285,219 | 1,280,285,219 | 100.00% |
| 250 Kb | 191 | 191 | 1,284,416,048 | 1,284,416,048 | 100.00% |
| 100 Kb | 214 | 214 | 1,287,435,802 | 1,287,435,802 | 100.00% |
| 50 Kb | 333 | 333 | 1,295,198,434 | 1,295,198,434 | 100.00% |
| 25 Kb | 333 | 333 | 1,295,198,434 | 1,295,198,434 | 100.00% |
| 10 Kb | 333 | 333 | 1,295,198,434 | 1,295,198,434 | 100.00% |
| 5 Kb | 333 | 333 | 1,295,198,434 | 1,295,198,434 | 100.00% |
| 2.5 Kb | 333 | 333 | 1,295,198,434 | 1,295,198,434 | 100.00% |
| 1 Kb | 333 | 333 | 1,295,198,434 | 1,295,198,434 | 100.00% |
| 0 bp | 333 | 333 | 1,295,198,434 | 1,295,198,434 | 100.00% |

**Screening and Final Assembly Releases**

Scaffolds that were not anchored in a chromosome were classified into bins depending on sequence content. Contamination was identified using blastn against the NCBI non-redundant nucleotide collection (NR/NT) and blastx using a set of known microbial proteins. Additional scaffolds were classified in the version 1.0 HAP1 release as redundant (unanchored scaffolds composed of >=95% 24mers >2x in all scaffolds) (138 scaffolds, 10.9 Mb), fungal (54 scaffolds, 3.2 Mb), repetitive (<=250 Kb scaffolds composed of >=95% 24mers >4x in >=5 Mb scaffolds) (21 scaffolds, 1.9 Mb), chloroplast (1 scaffolds, 158.8 Kb), and mitochondria (1 scaffolds, 21.3 Mb). Resulting final statistics for the HAP1 version 1.0 release are shown in Table S5.

**Table S5**. Final summary assembly statistics for the version 1.0 HAP1 chromosome scale assembly.

| **Scaffold total** | 12 |
| --- | --- |
| **Contig total** | 195 |
| **Scaffold sequence total** | 1,285.3 Mb |
| **Chromosome Sequence** | 1,283.3 Mb |
| **Contig sequence total** | 1,283.4 Mb (0.1% gap) |
| **Scaffold N/L50** | 5 / 134.5 Mb |
| **Contig N/L50** | 33 / 11.4 Mb |

Additional scaffolds were classified in the version 1.0 HAP2 release as redundant (unanchored scaffolds composed of >=95% 24mers >2x in all scaffolds) (39 scaffolds, 4.3 Mb), fungal (28 scaffold, 2.1 Mb), repetitive (<=250 Kb scaffolds composed of >=95% 24mers >4x in >=5 Mb scaffolds) (16 scaffolds, 1.5 Mb), chloroplast (1 scaffolds, 158.8 Kb), and mitochondria (1 scaffolds, 21.3 Kb). Resulting final statistics for the HAP2 version 1.0 release are shown in Table S6.

**Table S6**. Final summary assembly statistics for the version 1.0 HAP2 chromosome scale assembly.

| **Scaffold total** | 10 |
| --- | --- |
| **Contig total** | 169 |
| **Scaffold sequence total** | 1,274.7 Mb |
| **Chromosome Sequence** | 1,273.1 Mb |
| **Contig sequence total** | 1,273.1 Mb (0.1% gap) |
| **Scaffold N/L50** | 5 / 133.8 Mb |
| **Contig N/L50** | 32 / 13.7 Mb |
